# Interplay of macromolecular interactions during assembly of human DNA polymerase δ holoenzymes and initiation of DNA synthesis

**DOI:** 10.1101/2023.05.09.539896

**Authors:** Jessica L. Norris, Lindsey O. Rogers, Kara G. Pytko, Rachel L. Dannenberg, Samuel Perreault, Vikas Kaushik, Sahiti Kuppa, Edwin Antony, Mark Hedglin

## Abstract

In humans, DNA polymerase δ (Pol δ) holoenzymes, comprised of Pol δ and the processivity sliding clamp, proliferating cell nuclear antigen (PCNA), carry out DNA synthesis during lagging strand DNA replication, initiation of leading strand DNA replication, and the major DNA damage repair and tolerance pathways. Pol δ holoenzymes are assembled at primer/template (P/T) junctions and initiate DNA synthesis in a coordinated process involving the major single strand DNA-binding protein complex, replication protein A (RPA), the processivity sliding clamp loader, replication factor C (RFC), PCNA, and Pol δ. Each of these factors interact uniquely with a P/T junction and most directly engage one another. Currently, the interplay between these macromolecular interactions is largely unknown. In the present study, novel Förster Resonance Energy Transfer (FRET) assays reveal that dynamic interactions of RPA with a P/T junction during assembly of a Pol δ holoenzyme and initiation of DNA synthesis maintain RPA at a P/T junction and accommodate RFC, PCNA, and Pol δ, maximizing the efficiency of each process. Collectively, these studies significantly advance our understanding of human DNA replication and DNA repair.

---

In humans, DNA polymerase δ (Pol δ) holoenzymes, comprised of Pol δ and the processivity sliding clamp, proliferating cell nuclear antigen (PCNA), carry out DNA synthesis during lagging strand DNA replication, the initiation of leading strand DNA replication, and the major DNA damage repair and tolerance pathways^1–11^. In each of these DNA synthesis pathways, Pol δ holoenzymes are assembled at nascent primer/template (P/T) junctions and initiate DNA synthesis in a complex process that requires the spatially and temporally coordinated actions of many cellular factors that all interact with nascent P/T junctions; namely, the major single strand DNA (ssDNA)-binding protein complex, replication protein A (RPA), the processivity sliding clamp loader complex, replication factor C (RFC), PCNA, and Pol δ (**Figure 1A**)^12–22^. Each of these protein−DNA interactions with P/T junctions are unique (**Figure 1B**) and transpire in the context of the other protein−DNA interactions that assemble and disassemble over time. Furthermore, many of these protein-DNA interactions significantly overlap, implying mutual exclusivity and innate competition. Finally, nearly all of the aforementioned cellular factors (RPA, RFC, PCNA, Pol δ) directly engage one another through protein-protein interactions (**Figure 1C**). Currently, the interplay between the macromolecular interactions involved in the assembly of Pol δ holoenzymes and subsequent initiation of DNA synthesis is unknown, greatly limiting our fundamental understanding of human DNA replication and DNA damage repair. In the present study, we design and utilize a novel and efficient Förster Resonance Energy Transfer (FRET) assay that directly and continuously monitors the interactions of PCNA and RPA with P/T junctions as Pol δ holoenzymes are assembled and initiate DNA synthesis. Results reveal that dynamic interactions of RPA subunits with a P/T junction during assembly of a Pol δ holoenzyme and initiation of DNA synthesis both maintain RPA at the P/T junction and accommodate RFC, PCNA, and Pol δ, maximizing the efficiency of each process.

**Figure 1.**
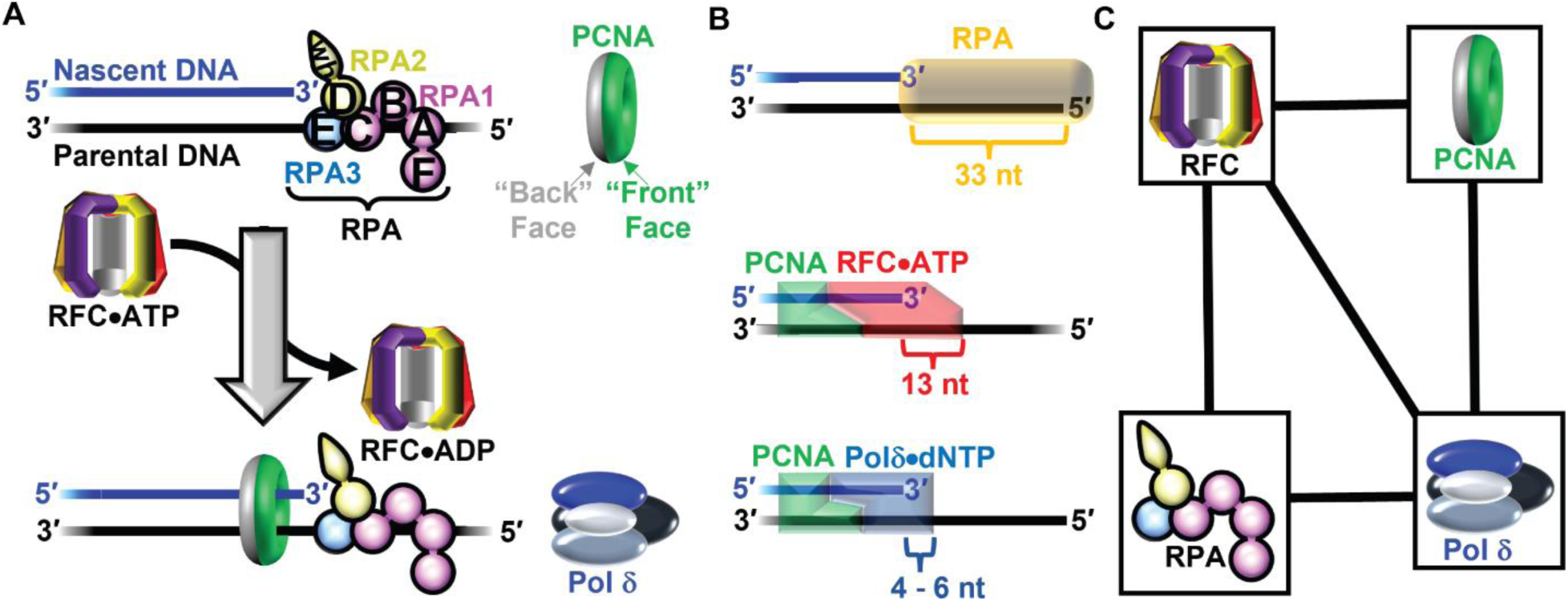
Assembly and macromolecular interactions of human Pol δ holoenzymes. (**A**) Assembly of human Pol δ holoenzymes. The subunits of human RPA are color-coded (RPA1 in pink, RPA2 in yellow, and RPA3 in blue) and depicted to illustrate the OB-folds (A – E) and winged-helix (wh) domain. A nascent P/T junction is pre-engaged by RPA in an orientation specific manner (as depicted). First, RFC utilizes ATP to load PCNA onto a nascent P/T junction in an orientation-specific manner such that the ‟front face” of the clamp is oriented towards the 3′ terminus of the primer from which DNA synthesis initiates. Next, Pol δ engages the ‟front face” of the PCNA sliding clamp encircling the nascent P/T junction, forming a holoenzyme, and subsequently initiates DNA synthesis (not shown). (**B**) Isolated protein•DNA interactions during assembly of the Pol δ holoenzyme. Shown in *Top*, *Middle*, and *Bottom* are the DNA footprints of the RPA complex, the activated RFC•ATP•‟Open” PCNA complex (i.e., activated loading complex), and the Pol δ holoenzyme, respectively. For each, the respective DNA region protected (i.e., DNA footprint) is depicted as a given protein complex (shown in color as a simple cartoon shape) overlaid on a P/T junction comprised of a 29 bp duplex region and a 33 nt ssDNA overhang. The protected ssDNA region is indicated for each. (**C**) Protein•protein interactions of the protein complexes involved in Pol δ holoenzyme assembly. Direct interactions are indicated by a solid black line connecting two respective protein complexes.

## Results

### RPA OBA maintains contacts with a P/T junction throughout loading of the resident PCNA

The ssDNA immediately adjacent to a nascent P/T junction is initially engaged by an RPA heterotrimer in an orientation-specific manner, as depicted in **Figure 1A**. In short, oligonucleotide binding (OB) fold A (OBA) of the RPA1 subunit aligns closer to the 5′ end of the ssDNA sequence and OBD of the RPA2 subunit aligns closer to the 3′ end such that OBD and OBC directly contact the 3′ terminus of the primer strand^12–19^. Next, RFC loads PCNA onto a nascent P/T junction that is engaged by RPA. Specifically, RFC, in complex with ATP, engages the “front face” of a free PCNA clamp in solution, opens the sliding clamp, and the RFC•ATP•‟Open” PCNA complex, referred to herein as the loading complex, engages a nascent P/T junction such that the “front face” of the PCNA clamp is oriented towards the 3′ terminus of the primer from which DNA synthesis initiates. Loading complexes may be targeted to P/T junctions engaged by RPA through direct protein•protein interactions between subunits 1 – 3 of the RFC complex and the RPA1 subunit of the RPA complex (**Figure 1C**)^23,24^. Upon engaging a nascent P/T junction, the loading complex adopts an activated conformation in which ATP hydrolysis by RFC is optimized. Herein, the activated conformation of the loading complex is referred to as the activated loading complex. ATP hydrolysis by RFC within the activated loading complex simultaneously closes PCNA around the DNA and the closed (i.e., loaded) PCNA is subsequently released onto the double stranded DNA (dsDNA) region of the nascent P/T junction. The resultant RFC•ADP complex then releases into solution, and this step is rate limited by dissociation of the RFC•ADP complex from the RPA1 subunit of the resident RPA ^20,21,23,25^. After completion of PCNA loading, the resident RPA stabilizes the loaded PCNA at/near the nascent P/T junction by prohibiting RFC-catalyzed unloading of PCNA and diffusion of PCNA along the adjacent ssDNA^25^.

RPA maintains contact(s) with a nascent P/T junction throughout loading of the resident PCNA^25,26^. Given the significant overlap in the DNA footprints of RPA and the activated complex on nascent P/T junctions (**Figure 1B**), it is unclear how this is possible. Specifically, both RPA and RFC (as part of the activated loading complex) directly engage the 3′ terminus of the primer strand and at least 13 nucleotides (nt) of the template ssDNA strand immediately downstream of a nascent P/T junction^12–19,24^. To address this long-standing enigma, we design and utilize novel FRET assays to directly and continuously monitor RPA-DNA interactions throughout the PCNA loading process described above. Initially, we focus on the interaction(s) of RPA OBA with nascent P/T junctions during association and activation of the loading complex.

To establish and monitor interactions of loading complexes with P/T junctions, we utilized a previously characterized Cy3, Cy5 FRET pair comprised of a Cy3-labeled P/T DNA substrate (5′ddPCy3/T, **Figure S1**) and Cy5-labeled PCNA (Cy5-PCNA) ^22,25,27^. The primer strand of the 5′ddPCy3/T DNA substrate contains a Cy3 donor near the 5′ end and is terminated at the 5′ end with biotin and at the 3′ end with a dideoxynucleotide (ddC). When pre-bound to NeutrAvidin, biotin attached to the 5′ end of a primer strand prevents loaded PCNA from diffusing off the dsDNA end of the substrate. For consistency, NeutrAvidin is included in all experiments in the present study that utilize 5′ biotin-terminated primers. A 3′ dideoxy-terminated primer cannot be extended by a DNA polymerase and is implemented here in consideration of subsequent experiments in the present study that utilize human DNA polymerase δ (pol δ). The Cy5-labeled PCNA homotrimer contains a single Cy5 acceptor at amino acid position 107 of a PCNA monomer. When encircling a P/T junction, either as part of a loading complex or as loaded PCNA, the Cy5 acceptor on the “back face” of the PCNA homotrimer is oriented towards the Cy3 donor on the 5′ddPCy3/T DNA substrate, yielding a FRET. Experiments are performed with adenosine 5′-O-(3-thio) triphosphate (ATPγS), an ATP analog in which one of the non-bridging oxygens of the γ-phosphoryl group is substituted for sulfur. This substitution severely inhibits hydrolysis of ATP by human RFC and, hence, stalls the PCNA loading pathway at activation of the loading complex^21,28^.

For a Cy3, Cy5 FRET pair, such as those described above, the gain and loss of FRET are directly indicated only by simultaneous, anti-correlated changes in the fluorescence emission intensities of the Cy3 donor and Cy5 acceptor. Specifically, appearance of (or increase in) FRET is indicated by a decrease (i.e., quenching) of Cy3 donor fluorescence emission intensity (*I*_563_) together with a concomitant increase in sensitized Cy5 acceptor fluorescence emission intensity (*I*_665_). Likewise, disappearance of (or decrease in) FRET is indicated by a decrease in sensitized *I*_665_ together with a concomitant increase (i.e., de-quenching) of *I*_563_. Therefore, to directly and continuously monitor change in FRET over time, we utilized a spectrofluorometer that monitors *I*_563_ and *I*_665_ essentially simultaneously (Δt = 0.235 ms) and permits successive additions of components to a given reaction mixture. The approximate FRET efficiency, E_FRET_, is then calculated for each *I*_563_, *I*_665_ pair over time. To analyze binding and activation of loading complexes on P/T junctions via FRET, the 5′ddPCy3/T DNA substrate is first pre-bound with NeutrAvidin and native RPA and I_563_ and *I*_665_ are monitored over time (**Figure 2A**). Then, loading complex pre-formed with RFC, Cy5-PCNA and ATPγS is added to the reaction mixture, the resultant solution is mixed, and the fluorescence emission intensity recording is resumed. Under the conditions of the assay, the loading complex rapidly engages the P/T junction and then the PCNA loading pathway stalls prior to ATP hydrolysis. Thus, an increase in E_FRET_ encompasses all kinetic steps along the PCNA loading pathway up to but not including hydrolysis of ATP by RFC within the activated loading complex^21,25,28^.

**Figure 2.**
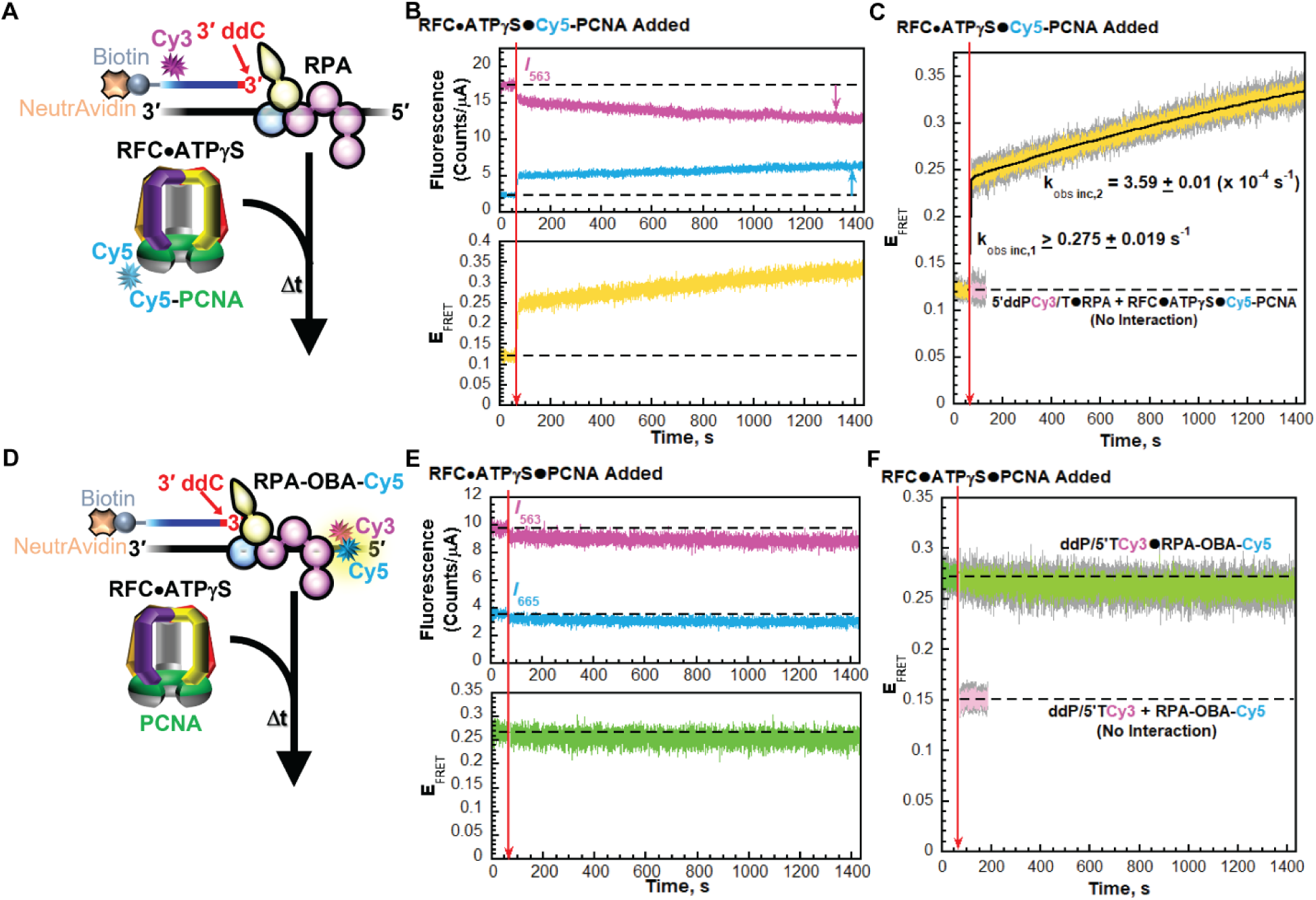
Interplay of RPA OBA and RFC interactions with nascent P/T junctions during PCNA loading in the presence of ATPγS. (**A - C**) Monitoring interactions of a loading complex with a P/T junction engaged by RPA. (**A**) Schematic representation of the FRET experiment performed with 5′ddPCy3/T (+ NeutrAvidin), RPA, and loading complex pre-formed with RFC, ATPγS, and Cy5-PCNA. (**B**) Sample time trajectories of *I*_563_ and *I*_665_ (*Top*) and their E_FRET_ (*Bottom*). The time at which the loading complex is added is indicated by a red arrow. Changes in *I*_563_ and *I*_665_ are indicated by magenta and cyan arrows, respectively. For observation, the *I*_563_, *I*_665_, and E_FRET_ values observed prior to the addition of the loading complex are fit to flat lines that are extrapolated to the axis limits. (**C**) FRET data. Each E_FRET_ trace is the mean of at least three independent traces with the S.E.M. shown in grey. The time at which the loading complex is added is indicated by a red arrow. The E_FRET_ trace observed after the addition of the loading complex is fit to a double exponential rise and the observed rate constants are reported in the graph. The predicted E_FRET_ trace (pink) for no interaction between 5′ddPCy3/T•RPA complex and the loading complex is fit to a flat line. (**D** – **F**) Monitoring RPA OBA interactions with a P/T junction that is engaged by a loading complex. (**D**) Schematic representation of the experiment. Reactions were carried out exactly as described for panel **A** above except that ddP/5′TCy3 DNA, RPA-OBA-Cy5, and PCNA were utilized. (**E**) Sample time trajectory of *I*_563_ and *I*_665_ (*Top*) and their E_FRET_ (*Bottom*) are plotted as described for panel **B** above. (**F**) FRET data. Each E_FRET_ trace is the mean of at least three independent traces with the S.E.M. shown in grey. Data is plotted as described for panel **C** above. The E_FRET_ trace observed prior to the addition of the loading complex is fit to a flat line that is extrapolated to the axis limits. The predicted E_FRET_ trace (pink) for no interaction between RPA-OBA-Cy5 and the ddP/5′TCy3 DNA is fit to a flat line.

Upon addition of the loading complex, *I*_665_ rapidly increases concomitantly with a rapid decrease in *I*_563_ after which *I*_665_ continues to increase slowly while *I*_563_ continues to decrease slowly (**Figure 2B**, *Top*). These synchronized, anti-correlated changes in *I*_563_ and *I*_665_ are indicative of the appearance and increase FRET (**Figure 2B**, *Bottom*). As observed in **Figure 2C**, E_FRET_ traces increase to values significantly above the E_FRET_ traces predicted for no interaction between the loading complex (with Cy5-PCNA and ATPγS) and 5′ddPCy3/T DNA. The increase in E_FRET_ is comprised of two phases (i.e., biphasic) with a lower limit for the observed rate constant of the first phase of *k*_obs inc,1_ ≥ 0.275 ± 0.019 s^-1^ and an observed rate constant of the second phase of *k*_obs inc,2_ = 3.59 ± 0.01 (x 10^-4^ s^-1^). A lower limit of *k*_obs inc,1_ is indicated due to the kinetic fitting of relatively few time points within its lifetime. The biphasic behavior and kinetic variables observed in the present study agree with that observed in a previous report that analyzed PCNA loading under similar conditions by monitoring sensitized Cy5 acceptor fluorescence emission intensity (*I*_665_) via stopped flow^25^. Next, we repeated these assays utilizing native PCNA and an alternative FRET pair to monitor interactions of RPA OBA with P/T junctions (**Figure 2D**). The Cy3-labeled P/T DNA substrate (ddP/5′TCy3 DNA, **Figure S1**) is identical to 5′ddPCy3/T DNA substrate described above except that the Cy3 donor is located towards the 5′ end of the template strand, rather than the 5′ end of the primer strand. The Cy5-labeled RPA (RPA-OBA-Cy5) contains a Cy5 acceptor on OBA of the RPA1 subunit that faces the Cy3 donor of ddP/5′TCy3 when engaged with the P/T junction (**Figure 2D**, **Figures S4 – S6**)^29^. RPA-OBA-Cy5 fully supports RFC-catalyzed loading of PCNA onto P/T junctions (**Figure S7** and **Table S1**).

The ddP/5′TCy3 DNA substrate is first pre-bound with NeutrAvidin and RPA-OBA-Cy5 and *I_563_* and *I*_665_ are monitored over time (**Figure 2D**). Here, a significant, constant E_FRET_ is observed prior to the addition of RFC in any form due to the stable interaction of RPA-OBA-Cy5 with ddP/5′TCy3 DNA (**Figures S4 – S6**). Next, loading complex pre-formed with RFC, native PCNA, and ATPγS is added, and the fluorescence emission intensities are monitored over time. Upon addition of the loading complex, both *I*_665_ and *I*_563_ rapidly and very slightly decrease then very slowly decrease over time (**Figure 2E**, *Top*). These synchronized, correlated changes in the fluorescence emission intensities are not attributable to FRET. Thus, any apparent change in the E_FRET_ values observed after the addition of the pre-formed loading complex is due to nonspecific effects^30^. For the example E_FRET_ trajectory depicted in **Figure 2E** (*Bottom*), the E_FRET_ values observed prior to the addition of the loading complex are maintained after addition of the pre formed loading complex. For the averaged E_FRET_ trajectory depicted in **Figure 2F**, the E_FRET_ traces observed prior to the addition of the loading complex persist and are maintained at the significantly elevated level above the E_FRET_ traces predicted for no interaction between ddP/5′TCy3 DNA and RPA-OBA-Cy5. This indicates that during binding and activation of the loading complex at a P/T junction the distance/orientation of RPA OBA relative to the 5′ end of the template strand does not change. Thus, the interaction(s) of RPA OBA with a P/T junction are not affected by interactions of activated complexes with the RPA1 subunit of RPA or any kinetic step along the PCNA loading pathway up to but not including hydrolysis of ATP by RFC within the activated loading complex. Next, we performed similar assays with ATP (**Figure 3**) to investigate the interaction(s) of RPA OBA with a P/T junction during the remainder of the PCNA loading pathway.

**Figure 3.**
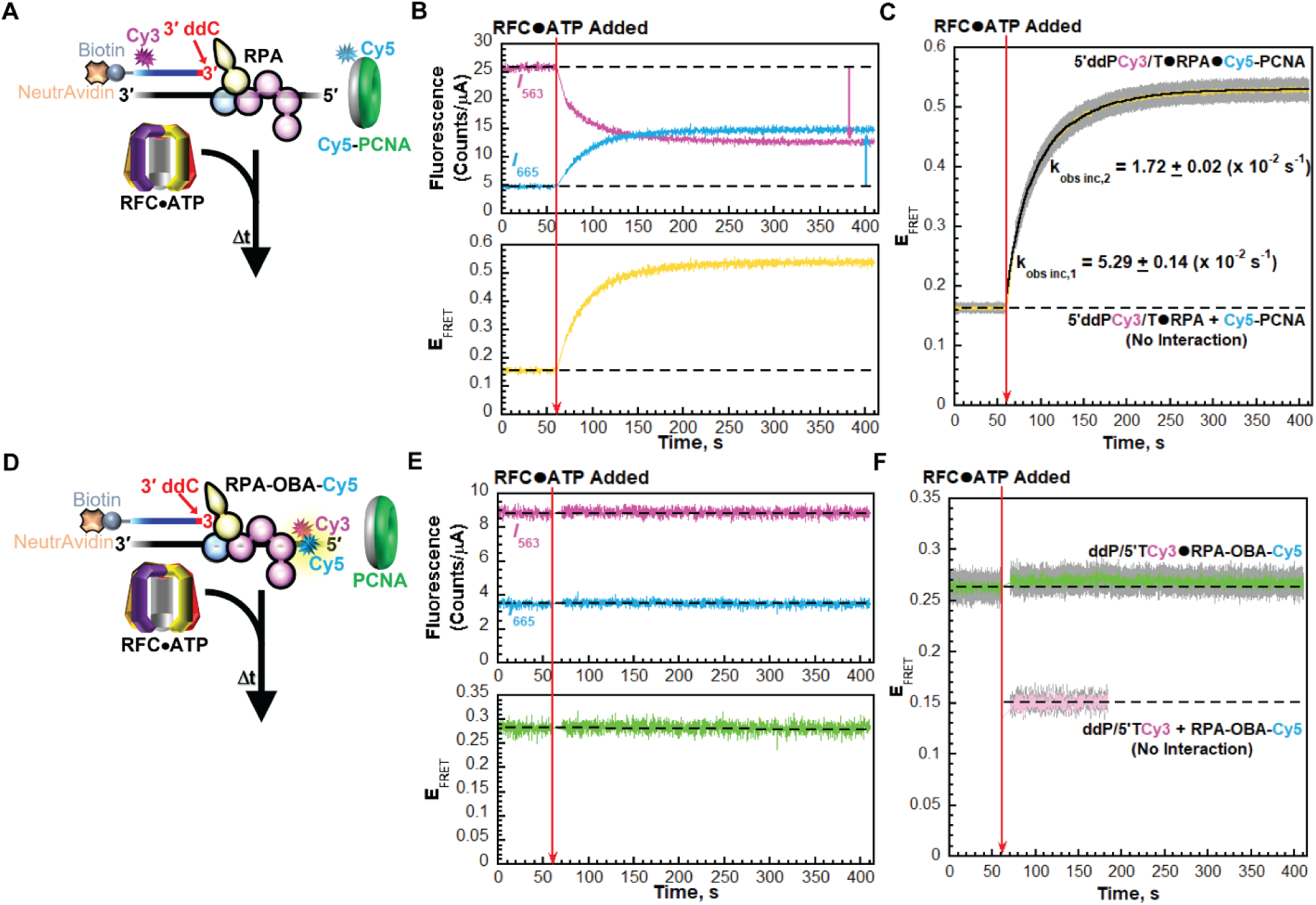
Interplay of RPA OBA and RFC interactions with nascent P/T junctions during PCNA loading in the presence of ATP. (**A - C**) Monitoring interactions of RFC and PCNA with a P/T junction engaged by RPA. (**A**) Schematic representation of the FRET experiment performed with 5′ddPCy3/T (+ NeutrAvidin), RPA, Cy5-PCNA and a pre-formed RFC•ATP complex. (**B**) Sample time trajectories of *I*_563_ and *I*_665_ (*Top*) and their E_FRET_ (*Bottom*). The time at which the RFC•ATP complex is added is indicated by a red arrow. Changes in *I*_563_ and *I*_665_ are indicated by magenta and cyan arrows, respectively. For observation, the *I*_563_, *I*_665_, and E_FRET_ values observed prior to the addition of the RFC•ATP complex are fit to flat lines that are extrapolated to the axis limits. (**C**) FRET data. Each E_FRET_ trace is the mean of at least three independent traces with the S.E.M. shown in grey. The time at which the RFC•ATP complex is added is indicated by a red arrow. The E_FRET_ trace observed prior to the addition of the RFC•ATP complex represents the complete absence of interactions between the 5′ddPCy3/T•RPA complex and Cy5-PCNA and is fit to a flat line that is extrapolated to the axis limits. The E_FRET_ trace observed after the addition of the RFC•ATP complex is fit to a double exponential rise and the observed rate constants are reported in the graph as well as in **Table S1**. (**D** – **F**) Monitoring RPA OBA interactions with a P/T junction that is engaged by RFC. (**D**) Schematic representation of the experiment. Reactions were carried out exactly as described in panel **A** except with ddP/5′TCy3 DNA, RPA-OBA-Cy5, and PCNA. (**E**) Sample time trajectory of *I*_563_ and *I*_665_ (*Top*) and their E_FRET_ (*Bottom*) are plotted as described in panel **B**. (**F**) FRET data. Each E_FRET_ trace is the mean of at least three independent traces with the S.E.M. shown in grey. Data is plotted as described in panel **C**. The E_FRET_ trace observed prior to the addition of the RFC•ATP complex is fit to a flat line that is extrapolated to the axis limits. The predicted E_FRET_ trace (pink) for no interaction between RPA-OBA-Cy5 and the ddP/5′TCy3 DNA is fit to a flat line.

First, interactions of RFC and PCNA with P/T junctions were monitored. The 5′ddPCy3/T DNA substrate is pre-bound with NeutrAvidin and native RPA. Then, Cy5-PCNA is added and *I_563_* and *I*_665_ are monitored over time (**Figure 3A**). Here, a low, constant E_FRET_ is observed prior to the addition of RFC in any form due to the presence of both the Cy3 donor and the Cy5 acceptor. E_FRET_ values observed during this period are a true experimental baseline signal representing the absence of any interactions between Cy5-PCNA and the 5′ddPCy3/T DNA•RPA complex. Next, pre-formed RFC•ATP complex is added, and the fluorescence emission intensities are monitored. Here, RFC•ATP must first bind a “free” PCNA in solution in order for loading to proceed. Under the conditions of the assay, PCNA loading is stoichiometric and biphasic^25^; first, all steps up to and including release of the closed PCNA ring on the P/T junction are rate-limited by a kinetic step along the PCNA loading pathway that occurs prior to and much slower than binding of the loading complex to the P/T junction; second, the loaded PCNA repositions relatively slowly on the P/T junction concurrent with release of the RFC•ADP complex into solution via its dissociation from the RPA1 subunit of the resident RPA engaged with the P/T junction. Thus, the appearance and increase in E_FRET_ towards maximal values encompasses all kinetic steps along the PCNA loading pathway and reflects; 1) a relatively fast kinetic step(s) (*k*_obs inc,1_) along the PCNA loading pathway that occurs prior to and much slower than binding of the loading complex to the P/T junction and; 2) a relatively slower release of the RFC•ADP complex into solution via its dissociation from the resident RPA engaged with the P/T junction (*k*_obs inc,2_)^25^.

Upon addition of RFC•ATP, the observed changes in *I_665_* and *I*_563_ are synchronized and anti correlated (**Figure 3B**, *Top*), indicating the appearance and increase in FRET (**Figure 3B**, *Bottom*). As observed in **Figure 3C**, E_FRET_ traces rapidly increase to values significantly above the E_FRET_ values observed for no interaction between Cy5-PCNA and the 5′ddPCy3/T•RPA complex. As expected, the rapid increase in E_FRET_ observed upon addition of RFC is biphasic with observed rate constants of *k*_obs inc,1_ = 5.29 ± 0.14 (x 10^-2^ s^-1^) and *k*_obs inc,2_ = 1.72 ± 0.02 (x 10^-2^ s^-1^) (**Table S1**). Importantly, *k*_obs inc_,_2_ agrees very well with the values reported in a previous study that analyzed PCNA loading under similar conditions by monitoring sensitized Cy5 acceptor fluorescence emission intensity (*I*_665_) via stopped flow. As discussed above, k_obs inc_,_2_ reports on release of the RFC•ADP complex into solution via its dissociation from the RPA1 subunit of the resident RPA engaged with the P/T junction. This kinetic step occurs throughout the time-dependent increase in E_FRET_ observed in **Figure 3C** and accounts for ∼30 – 60% of the E_FRET_ signal at any point in time, based on fits of the kinetic data.

Next, we repeated these assays utilizing ddP/5′TCy3 DNA substrate, RPA-OBA-Cy5, and PCNA (**Figure 3D**) to monitor the interaction(s) of RPA OBA with P/T junctions. Upon addition of RFC•ATP, neither *I*_665_ nor *I*_563_ change over time (**Figure 3E**, *Top*). These persistent *I*_563_ and *I*_665_ values indicate that the observed E_FRET_ values observed prior to the addition of the RFC•ATP complex do not change upon its subsequent addition (**Figure 3E**, *Bottom*). As observed in **Figure 3F**, the E_FRET_ traces observed prior to the addition of the RFC•ATP complex persist and are maintained at the significantly elevated level above the E_FRET_ traces predicted for no interaction between ddP/5′TCy3 DNA and RPA-OBA-Cy5. This indicates that the distance/orientation of RPA OBA relative to the 5′ end of the template strand does not change as the RFC•ADP complex engages and releases from the RPA1 subunit. Altogether, the results presented in **Figures 2** and **3** indicate that the distance/orientation of RPA OBA relative to the 5′ end of the template strand does not change throughout the PCNA loading pathway. Thus, the interaction(s) of RPA OBA with a nascent P/T junction is maintained throughout loading of the resident PCNA and unaffected by interactions of RFC complexes (loading complexes, RFC•ADP) with the P/T junction and the RPA1 subunit of the resident RPA. Next, we focused on the interaction(s) of RPA OBD with nascent P/T junctions during loading of the resident PCNA.

### RPA OBD is transiently displaced from the primer terminus of a P/T junction during loading of the resident PCNA

To establish and monitor interactions of loading complexes with P/T junctions, we utilized the aforementioned Cy5-PCNA and a Cy3-labeled P/T DNA substrate (5′PCy3/T, **Figure S1**) substrate in which the primer strand contains a Cy3 donor near the 5′ end and is also terminated at the 5′ end with biotin; the 3′ terminus of the primer is not modified. As expected, experiments performed with ATPγS (**Figure 4A** – **C**) yielded nearly identical results to those observed for the 5′ddPCy3/T DNA substrate under identical conditions (**Figure 2A** – **C**). Specifically, the increase in E_FRET_ is biphasic with a lower limit for the observed rate constant of the first phase of *k*_obs inc,1_ ≥ 0.336 ± 0.039 s^-1^ and an observed rate constant of the second phase of *k*_obs inc,2_ = 7.98 ± 0.01 (x 10^-4^ s^-1^) (**Figure 4C**). Next, we repeated these assays utilizing native PCNA and an alternative FRET pair to monitor interactions of RPA OBD with P/T junctions (**Figure 4D**). The Cy3-labeled P/T DNA substrate (3′PCy3/T DNA, **Figure S1**) is identical to the 5′PCy3/T DNA substrate described above except that the Cy3 donor is located at the 3′ terminus of the primer strand, rather than near 5′ end of the primer strand. The Cy5-labeled RPA (Cy5-OBD-RPA) contains a Cy5 acceptor on OBD of the RPA2 subunit that faces the Cy3 donor of the 3′PCy3/T when engaged with the P/T junction (**Supplemental Information, Figures S4 – S6**)^29^. RFC catalyzed loading of PCNA onto P/T junctions is fully supported by Cy5-OBD-RPA (**Figure S8, Table S1**) and on the 3′PCy3/TDNA substrate (**Figure S9** – **S10**, **Table S1**). The latter results agree with a recent report that revealed that *S. cerevisiae* RFC does not discriminate against nucleotides at the 3′ terminus of the primer strand during clamp loading^31^.

**Figure 4.**
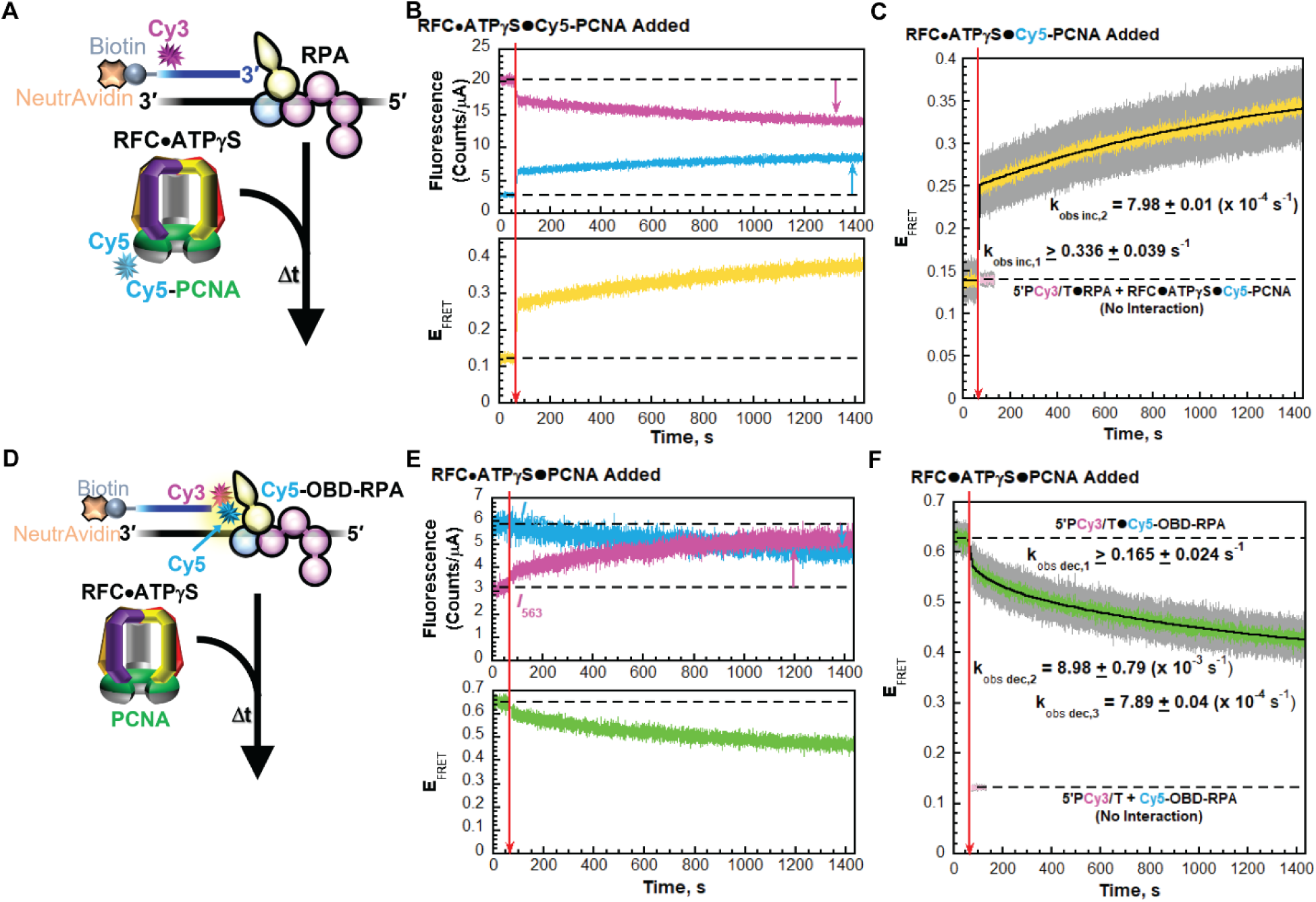
Interplay of RPA OBD and RFC interactions with nascent P/T junctions during PCNA loading in the presence of ATPγS. (**A - C**) Monitoring interactions of a loading complex with a P/T junction engaged by RPA. (**A**) Schematic representation of the FRET experiment performed with 5′PCy3/T (+ NeutrAvidin), RPA, and loading complex pre-formed with RFC, ATPγS, and Cy5-PCNA. (**B**) Sample time trajectories of *I*_563_ and *I*_665_ (*Top*) and their E_FRET_ (*Bottom*). The time at which the loading complex is added is indicated by a red arrow. Changes in *I*_563_ and *I*_665_ are indicated by magenta and cyan arrows, respectively. For observation, the *I*_563_, *I*_665_, and E_FRET_ values observed prior to the addition of the loading complex are fit to flat lines that are extrapolated to the axis limits. (**C**) FRET data. Each E_FRET_ trace is the mean of at least three independent traces with the S.E.M. shown in grey. The time at which the loading complex is added is indicated by a red arrow. The E_FRET_ trace observed after the addition of the loading complex is fit to a double exponential rise and the observed rate constants are reported in the graph. The predicted E_FRET_ trace (pink) for no interaction between 5P′Cy3/T•RPA complex and the loading complex is fit to a flat line. (**D** – **F**) Monitoring RPA OBD interactions with a P/T junction that is engaged by a loading complex. (**D**) Schematic representation of the experiment. Reactions were carried out exactly as described for panel **A** above except with P/5′TCy3 DNA, Cy5-OBD-RPA, and PCNA. (**E**) Sample time trajectory of *I*_563_ and *I*_665_ (*Top*) and their E_FRET_ (*Bottom*) are plotted as described for panel **B** above. (**F**) FRET data. Each E_FRET_ trace is the mean of at least three independent traces with the S.E.M. shown in grey. Data is plotted as described for panel **C** above. The E_FRET_ trace observed prior to the addition of the loading complex is fit to a flat line that is extrapolated to the axis limits. The predicted E_FRET_ trace (pink) for no interaction between Cy5-OBD-RPA and the P/5′TCy3 DNA is fit to a flat line.

The 3′PCy3/T DNA substrate is first pre-bound with NeutrAvidin and Cy5-OBD-RPA and *I_563_* and *I*_665_ are monitored over time (**Figure 4D**). A significant, constant E_FRET_ is observed prior to the addition of RFC in any form due to the stable interaction of Cy5-OBD-RPA with 3′PCy3/T DNA (**Figures S4 – S6**). Upon addition of the loading complex (RFC•ATPγS•Cy5 PCNA), *I*_665_ rapidly decreases concomitantly with a rapid increase in *I*_563_ after which *I*_665_ continues to decrease slowly while *I*_563_ continues to increase slowly (**Figure 4E**, *Top*). These synchronized, anti-correlated changes in *I*_563_ and *I*_665_ are indicative of a decrease in FRET (**Figure 4E**, *Bottom*). The decrease in E_FRET_ observed in **Figure 4F** is comprised of three phases with observed rate constants of *k*_obs inc,1_ ≥ 0.165 ± 0.024 s^-1^, *k*_obs inc,2_ = 8.98 ± 0.79 (x 10^-3^ s^-1^), and *k*_obs inc,3_ = 7.89 ± 0.04 (x 10^-4^ s^-1^). *k*_obs inc,3_, which accounts for 67.4 ± 2.50 % of the E_FRET_ decrease in **Figure 4F**, is nearly identical to *k*_obs,2_ for the E_FRET_ increase observed in **Figure 4C**.

Interestingly, and in contrast to those observed in **Figure 2C** and **2F**, the E_FRET_ behaviors observed in **Figures 4C** and **4F** with ATPγS are anti-correlated and occur with very similar kinetics; E_FRET_ between PCNA and DNA increases (**Figure 4C**), indicating binding of the loading complex to the P/T junction and adoption of the activated conformation; E_FRET_ between RPA OBD and DNA decreases, indicating that the distance between RPA OBD and the 3′ terminus of the primer strand increases. Together, this suggests that during PCNA loading, RPAOBD releases from the 3′ terminus of the primer strand to accommodate binding and activation of the loading complex at the nascent P/T junction. Next, we performed similar assays with ATP (**Figure 5**) to investigate the interaction(s) of RPA OBD with a P/T junction during the remainder of the PCNA loading pathway.

**Figure 5.**
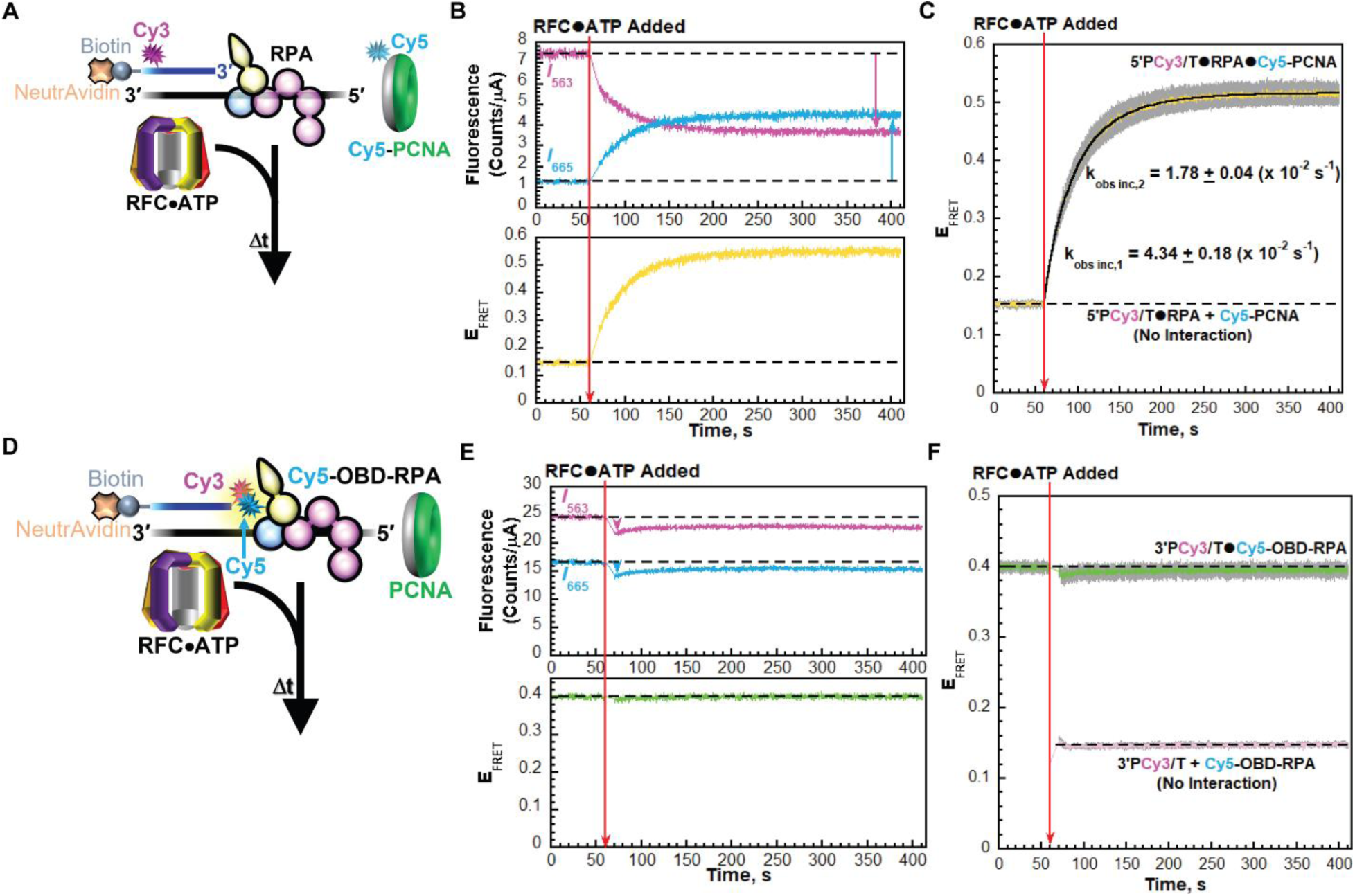
Interplay of RPA OBD and RFC interactions with nascent P/T junctions during PCNA loading in the presence of ATP. (**A - C**) Monitoring interactions of RFC and PCNA with a P/T junction engaged by RPA. (**A**) Schematic representation of the FRET experiment performed with 5′PCy3/T (+ NeutrAvidin), RPA, Cy5-PCNA and a pre-formed RFC•ATP complex. (**B**) Sample time trajectories of *I*_563_ and *I*_665_ (*Top*) and their E_FRET_ (*Bottom*). The time at which the RFC•ATP complex is added is indicated by a red arrow. Changes in *I*_563_ and *I*_665_ are indicated by magenta and cyan arrows, respectively. For observation, the *I*_563_, *I*_665_, and E_FRET_ values observed prior to the addition of the RFC•ATP complex are fit to flat lines that are extrapolated to the axis limits. (**C**) FRET data. Each E_FRET_ trace is the mean of at least three independent traces with the S.E.M. shown in grey. The time at which the RFC•ATP complex is added is indicated by a red arrow. The E_FRET_ trace observed prior to the addition of the RFC•ATP complex represents the complete absence of interactions between the 5′PCy3/T•RPA complex and Cy5-PCNA and is fit to flat line that is extrapolated to the axis limits. The E_FRET_ trace observed after the addition of the RFC•ATP complex is fit to a double exponential rise and the observed rate constants are reported in the graph as well as in **Table S1**. (**D** – **F**) Monitoring RPA OBD interactions with a P/T junction that is engaged by RFC. (**D**) Schematic representation of the experiment. Reactions were carried out exactly as described in panel **A** except with 3′PCy3/T DNA, Cy5-OBD-RPA, and PCNA. (**E**) Sample time trajectory of *I*_563_ and *I*_665_ (*Top*) and their E_FRET_ (*Bottom*) are plotted as described in panel **B**. (**F**) FRET data. Each E_FRET_ trace is the mean of at least three independent traces with the S.E.M. shown in grey. Data is plotted as described in panel **C**. The E_FRET_ trace observed prior to the addition of the RFC•ATP complex is fit to a flat line that is extrapolated to the axis limits. The predicted E_FRET_ trace (pink) for no interaction between Cy5-OBD-RPA and the 3′PCy3/T DNA is fit to a flat line.

First, interactions of RFC and PCNA with P/T junctions were monitored (**Figure 5A** – **C)**. The 5′PCy3/T DNA substrate is pre-bound with NeutrAvidin and native RPA. Then, Cy5-PCNA is added and *I*_563_ and *I*_665_ are monitored over time (**Figure 5A**). Here, a low, constant E_FRET_ is observed prior to the addition of RFC in any form due to the presence of both the Cy3 donor and the Cy5 acceptor. E_FRET_ values observed during this period represent a true experimental baseline signal representing the complete absence of interactions between Cy5-PCNA and the 5′PCy3/T DNA•RPA complex. Next, a pre-formed RFC•ATP complex is added, and the fluorescence emission intensities are monitored over time. Under the conditions of the assay, the appearance and increase in E_FRET_ towards maximal values encompasses all kinetic steps along the PCNA loading pathway and reflects; 1) a relatively fast kinetic step(s) (*k*_obs inc,1_) along the PCNA loading pathway that occurs prior to and much slower than binding of the loading complex to the P/T junction and; 2) a relatively slower release of the RFC•ADP complex into solution via its dissociation from the RPA1 subunit of the resident RPA complex engaged with the P/T junction (*k*_obs inc,2_)_25_.

Upon addition of RFC•ATP, the observed changes in *I*_665_ and *I*_563_ are synchronized and anti correlated (**Figure 5B**, *Top*), indicating the appearance and increase in FRET (**Figure 5B**, *Bottom*). As observed in **Figure 5C**, E_FRET_ traces rapidly increase to values significantly above the E_FRET_ values observed for no interaction between Cy5-PCNA and the 5′PCy3/T•RPA complex. As expected, the rapid increase in E_FRET_ observed upon addition of RFC is biphasic with observed rate constants of *k*_obs inc,1_ = 4.34 ± 0.18 (x 10^-2^ s^-1^) and *k*_obs inc,2_ = 1.78 ± 0.04 (x 10^-2^ s^-1^) (**Table S1**). Each of the observed rate constants agree very well with the values observed in **Figure 3C**. As discussed above, k_obs inc_,_2_ reports on release of the RFC•ADP complex into solution via its dissociation from the RPA1 subunit of the resident RPA engaged with the P/T junction. This kinetic step occurs throughout the time-dependent increase in E_FRET_ observed in **Figure 5C** and accounts for ∼30 – 60% of the E_FRET_ signal at any point in time, based on fits of the kinetic data. Release of the RFC•ADP complex into solution immediately follows hydrolysis of ATP by RFC within the activated loading complex (and concomitant closure of the sliding clamp ring) or release of the closed PCNA ring from the RFC•ADP complex^25^. Next, we repeated these assays utilizing the 3′PCy3/T DNA substrate, Cy5-OBD-RPA, and PCNA (**Figure 5D**) to monitor the interaction(s) of RPA OBD with P/T junctions.

A significant, constant E_FRET_ is observed prior to the addition of RFC in any form due to the stable interaction of Cy5-OBD-RPA with 3′PCy3/T DNA (**Figure 5E**, *Top,* **Figures S4 – S6**). After addition of RFC•ATP complex, RPA OBD subsequently releases from the 3′ terminus of the primer strand to accommodate binding and activation of the loading complex at the P/T junction (**Figure 4**). Under the conditions of the assay depicted in **Figure 5D**, kinetic steps along the PCNA loading pathway from binding of the loading complex to the P/T junction up to and including release of the closed PCNA ring on the DNA are kinetically invisible; only release of RFC•ADP into solution (via its dissociation from the RPA1 subunit of the resident RPA) is visible. This raises at least three scenarios for E_FRET_ traces observed after the addition of the RFC•ATP complex. First, if RPA OBD interactions are not subsequently re-established with the 3′terminus of the primer, E_FRET_ will instantaneously decrease upon addition of the RFC•ATP complex and then remain at a reduced level. Second, if RPA OBD interactions are re-established with the 3′ terminus of the primer prior to and much faster than release of RFC•ADP into solution, then a change in E_FRET_ will not be observed upon addition of the RFC•ATP complex. Third, if RPA OBD interactions are re-established concomitantly with or subsequent to release of RFC•ADP into solution, E_FRET_ will instantaneously decrease upon addition of the RFC•ATP complex and then increase over time to E_FRET_ values observed prior to the addition of the RFC•ATP complex.

Upon addition of RFC•ATP, the slight changes in *I*_665_ and *I*_563_ are synchronized and correlated (**Figure 5E**, *Top*). This behavior is due to nonspecific effects^30^. For the example E_FRET_ trajectory depicted in **Figure 5E** (*Bottom*), E_FRET_ values observed prior to the addition of the RFC•ATP complex are maintained after addition of the RFC•ATP loading complex. For the averaged E_FRET_ trajectory depicted in **Figure 5F**, the E_FRET_ traces observed prior to the addition of the RFC•ATP complex persist and are maintained at the significantly elevated level above the E_FRET_ traces predicted for no interaction between 3′PCy3/T DNA and Cy5-OBD-RPA. This agrees with the second scenario described above where RPA OBD interactions are re-established with the 3′ terminus of the primer prior to and much faster than release of RFC•ADP into solution via its dissociation from the RPA1 subunit of the resident RPA engaged with the P/T junction.

### Initiation of DNA synthesis by a Pol δ holoenzyme alters the orientation of PCNA encircling a P/T junction

In the next step of human Pol δ holoenzyme assembly, Pol δ engages the “front face” of PCNA encircling a P/T junction, forming a holoenzyme, and subsequently initiates DNA synthesis (**Figure 1A**). Currently, the interplay between the macromolecular interactions of Pol δ, PCNA, and RPA at nascent P/T junctions are unknown. In the present study, we utilize FRET assays to analyze these macromolecular interactions. Initially, we focus on interactions of PCNA with P/T junctions during formation of the Pol δ holoenzyme and subsequent initiation of DNA synthesis (**Figure 6A**).

**Figure 6.**
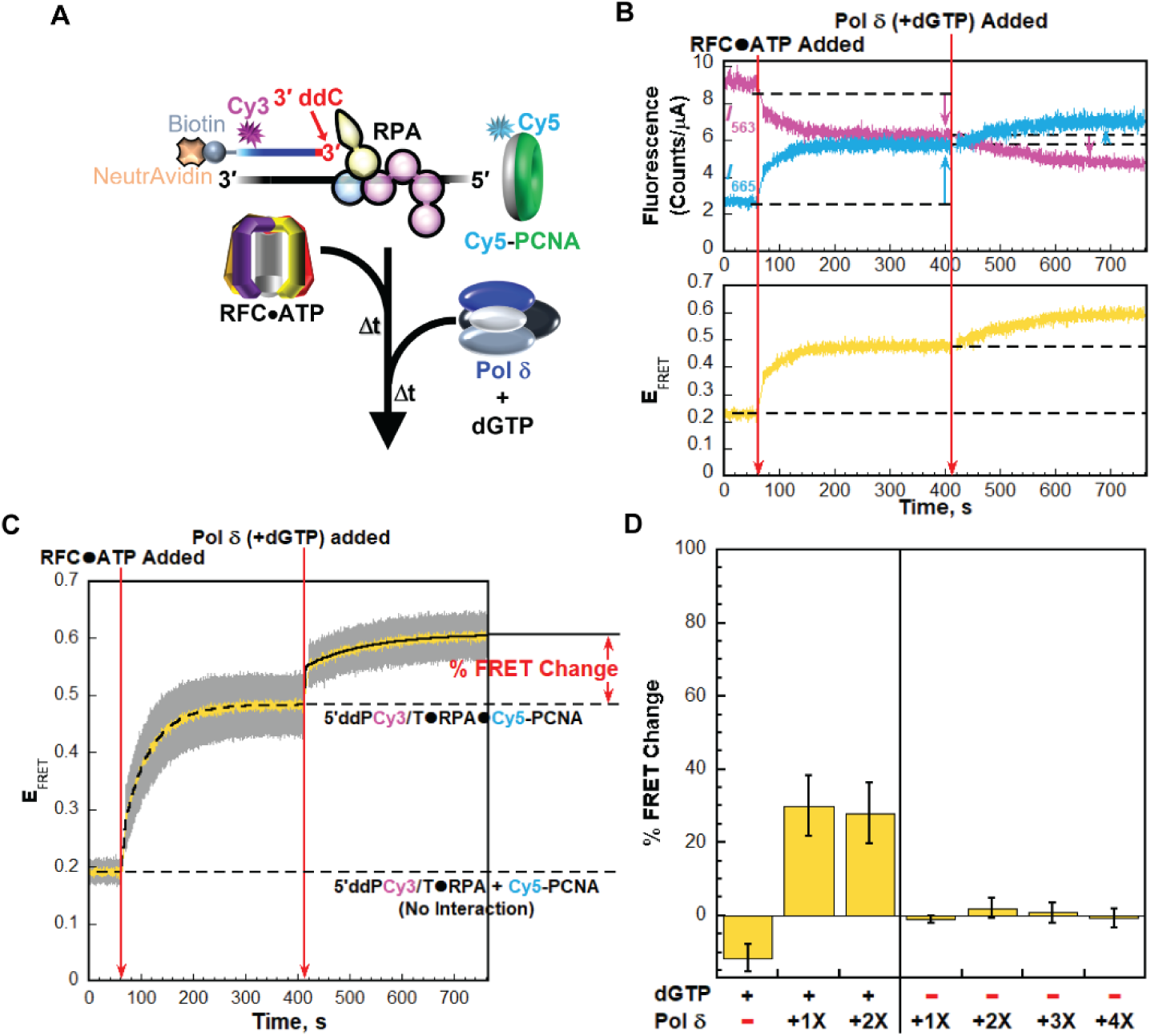
Formation of a Pol δ holoenzyme at a P/T junction engaged by RPA. (**A**) Schematic representation of the FRET experiment performed with 5′ddPCy3/T (+ NeutrAvidin), RPA, Cy5-PCNA, RFC, ATP, and Pol δ together with dGTP. (**B**) Sample time trajectories of *I*_563_ and *I*_665_ (*Top*) and their E_FRET_ (*Bottom*). The times at which the RFC•ATP complex and Pol δ (+ dGTP) are added are indicated by red arrows. For observation, the emission intensity traces and E_FRET_ values observed in the absence of RFC are each fit to dashed flat lines that are extrapolated. Also, dashed flat lines are drawn to highlight *I* values and E_FRET_ values observed at equilibrium for PCNA loading. Changes in *I*_563_ and *I*_665_ observed after each addition are indicated by magenta and cyan arrows, respectively. (**C)** FRET data. Each E_FRET_ trace is the mean of at least three independent traces with the S.E.M. shown in grey. The times at which the RFC•ATP complex and Pol δ (+ dGTP) are added are indicated by red arrows. The E_FRET_ trace observed prior to the addition of the RFC•ATP complex is fit to a dashed flat line that is extrapolated to the axis limits to depict the average E_FRET_ value for no interaction between Cy5-PCNA and the 5′ddPCy3/T•RPA complex. The E_FRET_ trace observed after the addition of the RFC•ATP complex is fit to a dashed double exponential rise that is extrapolated to the axis limits to depict the average E_FRET_ value for complete loading of Cy5-PCNA onto the Cy3-labeled P/T DNA substrate (i.e., the 5′ddPCy3/T•RPA•Cy5-PCNA complex). The E_FRET_ traces observed after the addition of Pol δ (+ dGTP) is fit to a double exponential rise. The % FRET Change observed after the addition of Pol δ (+ dGTP) is depicted in red. (**D**) Characterization of the % FRET Change observed. FRET experiments were repeated in the presence/absence of dGTP and with varying concentrations of Pol δ (0 – 100 nM heterotetramer) and the % FRET Change observed upon the ultimate addition (either dGTP, Pol δ + dGTP, or Pol δ) was measured. Each column is the mean of at least three independent replicates with the S.E.M. shown in black.

The 5′ddPCy3/T DNA substrate is pre-bound with NeutrAvidin and native RPA. Then, Cy5-PCNA is added, and *I_563_* and *I*_665_ are monitored over time. Next, pre-formed RFC•ATP is added, and the fluorescence emission intensities are monitored over time until PCNA loading is complete. Finally, dGTP is added together with Pol δ at a stoichiometric ratio of Pol δ to DNA and PCNA (i.e., DNA:PCNA:Pol δ = 1:1:1) and the fluorescence emission intensities are monitored over time. Under these conditions, Pol δ is stabilized in the initiation state for DNA synthesis where Pol δ engages PCNA encircling the P/T junction, the P/T junction, and aligns an incoming dGTP at the 3′ terminus of the primer strand in a correct base pair (bp) with the template nucleotide (C) immediately 5′ of the P/T junction^24,25,32^. Both extension of the primer (via Pol δ DNA polymerase activity) and degradation of the primer (via Pol δ exonuclease activity) are prohibited due to the utilization of a 3′ dideoxy-terminated primer and exonuclease deficient Pol δ^25^.

Upon addition of RFC•ATP, the observed changes in *I*_665_ and *I_563_* are synchronized and anti correlated (**Figure 6B**, *Top*), indicating the appearance and increase in FRET (**Figure 6B**, *Bottom*). As expected, E_FRET_ traces observed in **Figure 6C** rapidly increase in a biphasic manner upon addition of RFC•ATP and plateau at values significantly above the E_FRET_ traces observed for no interaction between Cy5-PCNA and 5′ddPCy3/T DNA. At the plateau, a PCNA is assembled onto each P/T junction and is rapidly and randomly diffusing along the dsDNA region (*D* = 2.24 x 10^7^ bp^2^/s)^33^. The biotin/NeutrAvidin complex at the 5′ terminus of the template strands prevent diffusion of loaded PCNA off the dsDNA end of the P/T DNA substrate. The resident RPA engaged at the P/T junction prohibits diffusion of PCNA along the adjacent ssDNA as well as RFC-catalyzed unloading of PCNA^25^. In the current experimental setup, rapid diffusion of loaded PCNA along the dsDNA region is kinetically invisible and, hence, E_FRET_ values observed at the plateau report on the average position of loaded PCNA relative to the 5′ end of the primer strand. Upon addition of stoichiometric Pol δ, *I*_665_ rapidly increases concomitantly with a rapid decrease in *I*_563_, after which both fluorescence emission intensities stabilize and persist over time (**Figure 6B**, *Top*). These synchronized, anti-correlated changes in *I*_563_ and *I*_665_ are indicative of a further increase in FRET (**Figure 6B**, *Bottom*). As observed in **Figure 6C**, upon addition of Pol δ, E_FRET_ traces rapidly increase to values significantly above the E_FRET_ values observed for the 5′ddPCy3/T•RPA•Cy5-PCNA complex (i.e., loaded Cy5-PCNA). The rapid increase in E_FRET_ observed upon addition of Pol δ is comprised of two phases (i.e., biphasic) with an observed rate constant for the slower phase (*k*_obs inc_,_2_) of 8.41 x 10^-3^ ± 0.19 (x 10^-3^ s^-1^). The overall E_FRET_ increase is due to a further reduction in the distance between the Cy3 donor near the 5′ end of the primer strand and the Cy5 acceptor on the “back” face of the loaded PCNA ring. Next, we further characterized the enhanced FRET state observed upon addition of Pol δ by monitoring the ‟% FRET Change” (depicted in **Figure 6C**) under various conditions (**Figure 6D**).

For reference, the % FRET change observed in **Figure 6C** is reported in **Figure 6D** (+ dGTP, 1X Pol δ). Doubling the concentration of Pol δ in the presence of dGTP (+ dGTP, 2X Pol δ) did not affect the % FRET Change, indicating that the enhanced FRET state is saturated at stoichiometric Pol δ. Next, dGTP was omitted. Under these conditions, Pol δ saturates PCNA encircling P/T junctions to form Pol δ holoenzymes but engages P/T junctions with dramatically low affinity, if at all^21,22,24,27,34,35^. Inclusion of only Pol δ (- dGTP, + 1X Pol δ) did not yield a % FRET Change and this behavior persisted at increasing concentrations of Pol δ (up to a DNA:PCNA:Pol δ ratio of 1:1:4). This indicates that the enhanced FRET state requires dGTP. Furthermore, these results reveal that Pol δ alone does not displace RPA from a P/T junction upon engaging the resident PCNA, as follows. A large protein, such as Pol δ, simply binding to PCNA encircling a P/T junction decreases the diffusion constant only 2.1-fold^33^. Thus, in the absence of dNTPs, if Pol δ displaced RPA from the 5′ddPCy3/T DNA upon engaging the resident Cy5-PCNA, the resultant Cy5-PCNA•Pol δ complex would rapidly diffuse off the ssDNA end of the 5′ddPCy3/T DNA simultaneously with RPA displacement. This would lead to a rapid decrease in E_FRET_ (rate-limited by displacement of RPA) and a subsequent, relatively slow increase in E_FRET_ as free Cy5-PCNA is reloaded by RFC. However, a % FRET Change is not observed in **Figure 6D** when only Pol δ is included, indicating that loaded PCNA persists on P/T DNA throughout Pol δ holoenzyme formation and, hence, RPA maintains contact(s) with the respective P/T junction throughout Pol δ holoenzyme formation. This is directly investigated in **Figure 7** below.

**Figure 7.**
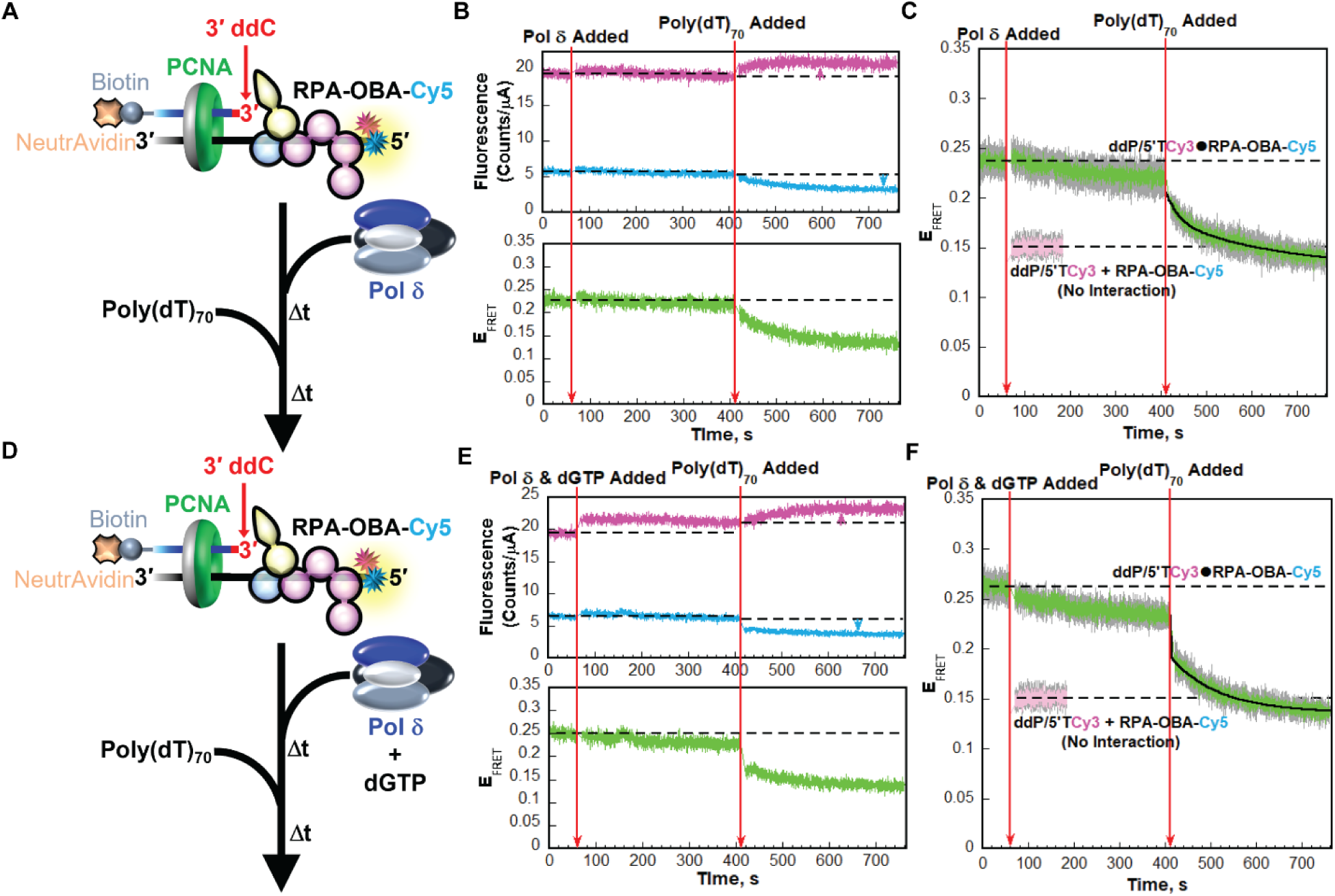
RPA dynamics during formation of a Pol δ holoenzyme and initiation of DNA synthesis. (**A**) Schematic representation of the FRET experiment performed with 5′ddP/5′TCy3 (+ NeutrAvidin), RPA-OBA-Cy5, PCNA, RFC, ATP, Pol δ, and poly(dT)_70_. (**B**) Sample time trajectories of *I*_563_ and *I*_665_ (*Top*) and their E_FRET_ (*Bottom*). The times at which Pol δ and poly(dT)_70_ are added are indicated by red arrows. For observation, the emission intensity traces and E_FRET_ values observed in the absence of Pol δ are each fit to dashed flat lines that are extrapolated. Also, dashed flat lines are drawn to highlight the *I* values and E_FRET_ values observed at equilibrium after the addition of Pol δ. Changes in *I*_563_ and *I*_665_ observed after the addition of poly(dT)_70_ are indicated by magenta and cyan arrows, respectively. (**C)** FRET data. Each E_FRET_ trace is the mean of at least three independent traces with the S.E.M. shown in grey. The times at which the Pol δ and poly(dT)_70_ are added are indicated by red arrows. The E_FRET_ trace observed prior to the addition of the Pol δ is fit to a dashed flat line that is extrapolated to the axis limits to depict the average E_FRET_ value for the interaction between RPA-OBA-Cy5 and ddP/5′TCy3. The predicted E_FRET_ trace (pink) for no interaction between RPA-OBA-Cy5 and the ddP/5′TCy3 DNA is fit to a flat line that is extrapolated to the axis limits. The E_FRET_ trace observed after the addition of the poly(dT)_70_ is fit to a double exponential decay. (**D**) Schematic representation of the FRET experiment performed in the same manner as that depicted in panel **A** except that Pol δ is added simultaneously with dGTP. (**E**) Sample time trajectories of *I*_563_ and *I*_665_ (*Top*) and their E_FRET_ (*Bottom*) plotted as in panel **B**. (**F)** FRET data plotted and analyzed as in panel **C**. Each E_FRET_ trace is the mean of at least three independent traces with the S.E.M. shown in grey.

Inclusion of only dGTP (+ dGTP, - Pol δ) led to a slight decrease in FRET due to the inhibition of PCNA re-loading by RFC (**Figure S11 - S13**)^36^. Thus, the enhanced FRET state requires Pol δ. Altogether, the results in **Figure 6D** indicate that formation of the enhanced FRET state following RFC-catalyzed loading of PCNA requires both dGTP and Pol δ. Hence, the enhanced FRET state represents the initiation state for DNA synthesis. Furthermore, these results reveal that formation of the initiation state for DNA synthesis alters the orientation/distance of the closed PCNA ring relative to the P/T junction. Next, we investigated the interactions of RPA with a P/T junction during formation of a Pol δ holoenzyme and subsequent initiation of DNA synthesis.

### RPA remains engaged with a P/T junction during formation of a Pol δ holoenzyme and initiation of DNA synthesis

Human Pol δ interacts (albeit with unknown affinity) with the RPA1 subunit of an RPA complex that is engaged with a P/T junction (**Figure 1C**) and, hence, these interactions may initially target Pol δ to P/T junctions where it captures a diffusing, loaded PCNA^23,37^. However, during initiation of DNA synthesis from a nascent P/T junction, the DNA footprints of RPA and Pol δ on a P/T junction significantly overlap (**Figure 1B**)^12–18,24,32^. To investigate the interplay between the macromolecular interactions of Pol δ and RPA at a nascent P/T junction, we performed FRET assays similar to those described above in **Figure 6** to monitor interactions of RPA OBA with a P/T junction during formation of a Pol δ holoenzyme and initiation of DNA synthesis (**Figure 7**).

The ddP/5′TCy3 DNA substrate is pre-bound with NeutrAvidin and RPA-OBA-Cy5 (**Figure 7A**). Then, PCNA is added followed by pre-formed RFC•ATP complex. After completion of RFC-catalyzed loading of PCNA, *I_563_* and *I*_665_ are monitored over time. Here, a significant, constant E_FRET_ is observed prior to the addition of Pol δ due to the stable interaction of RPA- OBA-Cy5 with ddP/5′TCy3 DNA (**Figures 2**, **3**, and **S4 – S6**). Next, Pol δ alone is added at a 2 fold excess to DNA and PCNA to ensure all loaded PCNA is engaged in a Pol δ holoenzyme and the fluorescence emission intensities are monitored over time. Finally, to demonstrate RPA occupancy of the P/T junction following Pol δ holoenzyme formation, a large excess of poly(dT)_70_ ssDNA is added, and the fluorescence emission intensities are monitored over time. Human RPA has exceptionally high affinity for ssDNA at physiological ionic strength but can undergo facilitated exchange due to the dynamic ssDNA-binding interactions of its individual OB folds, enabling RPA to exchange between ssDNA sequences when free ssDNA is present in solution (**Figure S14**)^29,38–40^. The ssDNA binding affinity of human RPA is highest for poly(dT) and each poly(dT)_70_ accommodates at least two RPA complexes^12–14,41^. Hence, poly(dT)_70_ serves as an effective trap to release RPA-OBA-Cy5 from the ddP/5′TCy3 DNA substrate via facilitated exchange and prohibit re-binding.

Upon addition of Pol δ alone, the slight changes in *I*_665_ and *I*_563_ are synchronized and correlated (**Figure 7B**, *Top*). For the example E_FRET_ trajectory depicted in **Figure 7B** (*Bottom*), the E_FRET_ values observed after addition of Pol δ increase very slightly and then slowly and minimally decrease over time, mirroring the synchronized, correlated changes observed for *I*_665_ and *I*_563_ over time (**Figure 7B**, *Top*). This apparent time-dependent change in the E_FRET_ values observed after the addition of Pol δ is due to nonspecific effects and not attributable to changes in the distance between the cyanine labels^30^. Upon addition of poly(dT)_70_, the observed changes in *I*_665_ and *I*_563_ are synchronized and anti-correlated (**Figure 7B**, *Top*), indicating a decrease in FRET (**Figure 7B**, *Bottom*). For the averaged E_FRET_ trajectory in **Figure 7C**, the E_FRET_ values observed prior to the addition of Pol δ increase very slightly after addition of Pol δ, and then slowly and minimally decrease. These time-dependent changes in E_FRET_ are due to indirect effects on the fluorescence emission intensities of Cy3 and Cy5 (**Figure 7B**, *Top*). Regardless, E_FRET_ values observed after addition of Pol δ remain within experimental error of the E_FRET_ values observed prior to the addition of Pol δ and also significantly elevated above the E_FRET_ trace predicted for no interaction between ddP/5′TCy3 DNA and Cy5-OBD-RPA. Upon addition of excess poly(dT)_70_, E_FRET_ rapidly decreases over time to the E_FRET_ trace predicted for no interaction between ddP/5′TCy3 DNA and RPA-OBA-Cy5, indicating release of RPA-OBA-Cy5 from ddP/5′TCy3 via facilitated exchange with the ssDNA trap. Together, this suggests that nearly all (≥ 85.1 ± 11.6%, based on the observed E_FRET_ changes), if not all, RPA maintains contact(s) with a P/T junction throughout formation of the resident Pol δ holoenzyme, which may include direct protein•protein interactions with Pol δ. This agrees with the results discussed above for **Figure 6D**.

Next, we repeated the experiments described in **Figure 7A** – **C** by adding dGTP simultaneously with a 2-fold excess of Pol δ compared to DNA and PCNA (**Figure 7D** – **F**). Here, excess polymerase ensures all loaded PCNA is engaged with Pol δ in the initiation state for DNA synthesis (**Figure 6D**). Upon addition of Pol δ and dGTP, the slight changes in *I*_665_ and *I*_563_ are synchronized and correlated (**Figure 7E**, *Top*). This behavior is due to nonspecific effects^30^. For the example E_FRET_ trajectory depicted in **Figures 7E** (*Bottom*) and the averaged E_FRET_ trajectory depicted in **Figures 7F**, the E_FRET_ values observed prior to the addition of Pol δ and dGTP slowly and minimally decrease after the addition of Pol δ and dGTP due to indirect effects on the fluorescence emission intensities of Cy3 and Cy5 (**Figure 7E**, *Top*). E_FRET_ values observed after addition of Pol δ and dGTP remain within experimental error of the E_FRET_ values observed prior to the addition of Pol δ and dGTP and also significantly elevated above the E_FRET_ trace predicted for no interaction between ddP/5′TCy3 DNA and Cy5-OBD-RPA. Upon addition of excess poly(dT)_70_, E_FRET_ rapidly decreases over time to the E_FRET_ trace predicted for no interaction between ddP/5′TCy3 DNA and RPA-OBA-Cy5, indicating release of RPA-OBA-Cy5 from ddP/5′TCy3 via facilitated exchange with the ssDNA trap. Together, this indicates that nearly all (≥ 77.3 ± 5.7% based on the observed changes in E_FRET_), if not all, RPA maintains contact(s) with a P/T junction during initiation of DNA synthesis by the resident Pol δ holoenzyme.

## Discussion

Human Pol δ holoenzymes are assembled at primer/template (P/T) junctions and initiate DNA synthesis in a complex process that requires the spatially and temporally coordinated actions of RPA, RFC, PCNA, and Pol δ. Each of these factors interact uniquely with a P/T junction and most directly engage one another. Currently, the interplay between these macromolecular interactions during Pol δ holoenzyme assembly and initiation of DNA synthesis is largely unknown. In the present study, we designed and utilized novel FRET assays to monitor these macromolecular interactions using recombinant human proteins. Together with previous work from our lab and others, the results from the present study provide the first complete description of human Pol δ holoenzyme assembly and initiation of DNA synthesis (**Figure 8**). The ssDNA immediately adjacent to a nascent P/T junction is initially engaged by an RPA heterotrimer in an orientation-specific manner (**Figure 1**). Specifically, 30 – 33 nt of ssDNA are engaged with a defined 5′→3′ polarity where OBA of the RPA1 subunit aligns closer to the 5′ end of the ssDNA sequence and OBD of the RPA2 subunit aligns closer to the 3′ end such that OBD and OBC directly contact the 3′ terminus of the primer strand. RPA remains engaged with the P/T DNA with exceptionally high affinity but exists in microscopically dissociated states due to the dynamic ssDNA-binding interactions of its individual OB folds. In particular, both OBA and OBD have been directly observed to rapidly bind to and dissociate from ssDNA while RPA remains engaged with ssDNA^12–19^. Next, RFC, in complex with ATP, engages the “front face” of a free PCNA in solution, opens the sliding clamp, and the resultant loading complex engages a nascent P/T junction such that the “front face” of PCNA is oriented towards the 3′ terminus of the primer from which DNA synthesis initiates (**Figure 8**, *Step 1*). Loading complexes may be targeted to P/T junctions engaged by RPA through direct protein•protein interactions between subunits 1 – 3 of the RFC complex and the RPA1 subunit of the RPA complex (**Figure 1C**)^23,24^. Such targeting, if it occurs, does not observably affect the interaction(s) of RPA OBA with a P/T junction (**Figures 2D** - **F**). Upon engaging a nascent P/T junction, the loading complex adopts an activated conformation in which ATP hydrolysis by RFC is optimized (**Figures 2A** - **C** and **4A** - **C**). During association and activation of the loading complex, RPA OBD is released from the 3′ terminus of the primer strand to accommodate binding and activation of the loading complex at the nascent P/T junction (**Figure 4D** – **F**) but RPA remains engaged with the P/T DNA through the persistent interaction(s) of at least RPA OBA with the template strand downstream of the P/T junction (**Figure 3D** – **F**). Given the dynamics of RPA•ssDNA complexes discussed above, we posit that a loading complex captures a P/T junction that is exposed during microscopic dissociation of RPA OBD, and likely RPA OBC, from the 3′ terminus of the primer strand and the template ssDNA immediately 5′ of the P/T junction. Microscopic dissociation of RPA OBA and other OB folds downstream of the nascent P/T junction do not manifest to microscopic dissociation of RPA.

**Figure 8.**
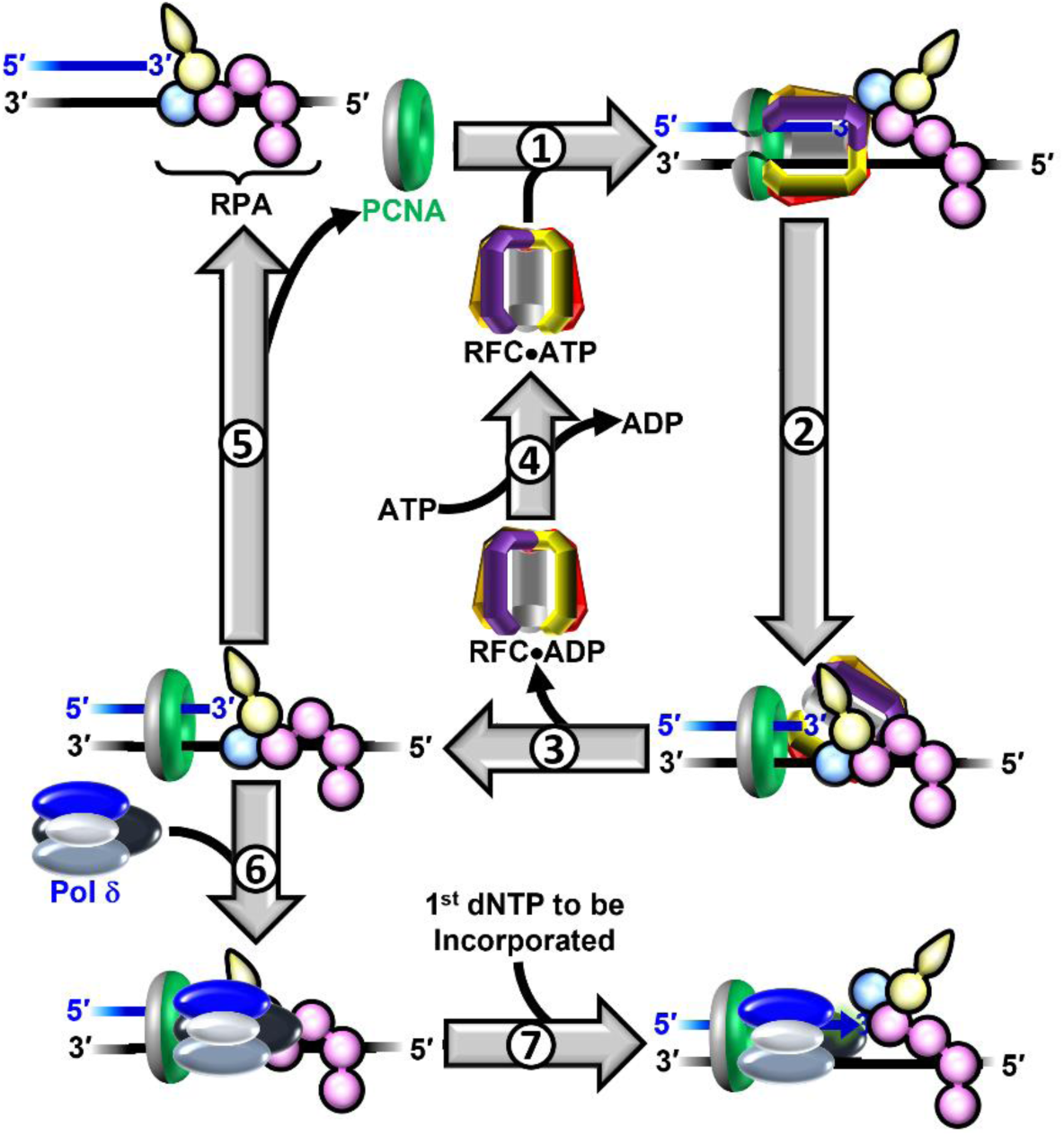
Assembly of the human Pol δ holoenzyme and initiation of DNA synthesis. A nascent P/T junction is pre-engaged by RPA in an orientation-specific manner (as depicted) and PCNA resides in solution (*Top Left*). **1**) RFC•ATP engages the ‟front face” of a free PCNA in solution, opens the sliding clamp, and the resultant loading complex engages a nascent P/T junction such that the ‟front face” of PCNA is oriented towards the 3′ terminus of the primer from which DNA synthesis initiates. Upon engaging a nascent P/T junction, the loading complex adopts an activated conformation in which ATP hydrolysis by RFC is optimized. RPA OBD is released from the 3′ terminus of the primer strand to accommodate binding and activation of the loading complex at the nascent P/T junction. **2**) ATP hydrolysis by RFC within the activated loading complex simultaneously closes PCNA around the DNA and the closed (i.e., loaded) PCNA is subsequently released onto the dsDNA region of the nascent P/T junction. Concomitant with closure of PCNA or release of loaded PCNA, the resultant RFC•ADP complex vacates the P/T junction, transferring to the RPA1 subunit, and the interactions of RPA OBD are re-established with the 3′ terminus of the primer strand. **3**) The RFC•ADP complex releases into solution via dissociation from the RPA1 subunit of the resident RPA engaged with the nascent P/T junction. **4**) RFC then exchanges ADP for ATP. **5**) In the absence of catalyzed unloading of PCNA and significant translocation of loaded PCNA, the only pathway for dissociation of Pol δ, loaded PCNA may dissociate from a nascent P/T junction via spontaneous opening of the PCNA ring. **6**) In the presence of Pol δ, the DNA polymerase engages the ‟front face” of loaded PCNA encircling a P/T junction, forming a holoenzyme. **7**) An assembled Pol δ holoenzyme engages a nascent P/T junction and an incoming dNTP and aligns the incoming dNTP at the 3′ terminus of the primer strand in a correct base pair with the template nt immediately 5′ of the P/T junction.

ATP hydrolysis by RFC within the activated loading complex simultaneously closes PCNA around the DNA and the closed (i.e., loaded) PCNA is subsequently released onto the dsDNA region of the nascent P/T junction (**Figure 8**, *Step 2*). RPA engaged with a nascent P/T junction promotes release of closed PCNA from RFC•ADP either directly or indirectly via stimulation of ATP hydrolysis by RFC within the activated loading complex (and simultaneous closure of the PCNA clamp)^25^. Concomitant with closure of PCNA or release of loaded PCNA, the resultant RFC•ADP complex vacates the P/T junction, transferring to the RPA1 subunit, and the interactions of RPA OBD are re-established with the 3′ terminus of the primer strand (**Figure 5D** - **F**). The RFC•ADP complex subsequently releases into solution (**Figure 8**, *Step 3*) and this step is rate-limited by dissociation of the RFC•ADP complex from the RPA1 subunit of the resident RPA engaged with the nascent P/T junction (**Figures 3A** – **C** and **5A** – **C**)^20,21,23,25^. RFC then exchanges ADP for ATP (**Figure 8**, *Step 4*) and is once again competent for PCNA loading.

Loaded PCNA rapidly and randomly diffuses (*D* = 2.24 x 10^7^ bp^2^/s) along the dsDNA region immediately 5′ of the P/T junction^33^. Diffusion of loaded PCNA away from a P/T junction is restricted by physical blocks, i.e., “protein roadblocks,” such as sub-nucleosomes and high affinity transcription factors that rapidly re-assemble on or rebind to nascent DNA generated during DNA replication^42–44^. This is mimicked in the current experimental setup where biotin/NeutrAvidin complexes at the 5′ termini of the template strands prevent diffusion of loaded PCNA off the dsDNA end of the P/T DNA substrates. The resident RPA engaged at the P/T junction prohibits diffusion of PCNA along the adjacent ssDNA as well as RFC-catalyzed unloading of PCNA^25^. Recent *in vivo* evidence suggests that enzyme-catalyzed unloading of PCNA from a P/T junction will not occur until the primer is completely extended and ligated to the downstream duplex region^45^. In the absence of catalyzed unloading of PCNA and significant translocation of loaded PCNA, the only pathway for dissociation of loaded PCNA from a nascent P/T junction is through spontaneous opening of the PCNA ring, which is dramatically slow [*k*_open_ = 1.25 ± 0.32 (x 10^-3^) s^−1^]^46^, ∽20-fold slower than the observed rate constant RFC-catalyzed loading of “free” PCNA onto a P/T junction in the presence of ATP (**Figure 3, 5, S3, S3 – S9**, and **Table 1**). Thus, upon dissociation of PCNA from a nascent P/T junction via spontaneous opening of the PCNA ring (**Figure 8**, *Step 5*), RFC utilizes ATP to instantly reload PCNA back onto the nascent P/T junction (**Figure 8**, *Steps 1* - *3*) such that the loss of loaded PCNA from a nascent P/T junction is very transient and essentially not observed (**Figure S11**)^21,25^.

In the next step of Pol δ holoenzyme assembly, Pol δ engages the “front face” of loaded PCNA encircling a P/T junction, forming a holoenzyme (**Figure 8**, *Step 6*). Human Pol δ is comprised of four subunits, three of which contain PCNA-binding motifs and simultaneously bind all subunits within a given PCNA homotrimer. This multivalent interaction leads to a significantly high affinity for PCNA encircling P/T junctions (K_D_ < 10 nM)^34^ . In the absence of dNTPs (i.e., no DNA synthesis), Pol δ has dramatically low affinity, if any, for P/T junctions ^22,24,25,27,34^. Thus, it is likely that Pol δ directly engages a loaded PCNA that is rapidly and randomly diffusing along the dsDNA region of a nascent P/T junction. However, human Pol δ interacts (albeit with unknown affinity) with the RPA1 subunit of an RPA complex that is engaged with a P/T junction (**Figure 1C**). Thus, it is possible that these interactions initially target Pol δ to a nascent P/T junction where it captures a diffusing, loaded PCNA^23,37^. Such targeting, if it occurs, as well as formation of a Pol δ holoenzyme do not displace the resident RPA from nascent P/T junction nor observably affect the interaction(s) of RPA OBA with a P/T junction (**Figure 6D** and **7A- C**).

Finally, an assembled Pol δ holoenzyme engages a nascent P/T junction and an incoming dNTP and aligns the incoming dNTP at the 3′ terminus of the primer strand in a correct bp with the template nucleotide immediately 5′ of the P/T junction (**Figure 8**, *Step 7*)^24,25,32^. The results presented in **Figure 6** indicate that formation of the initiation state for DNA synthesis alters the orientation/distance of the closed PCNA ring relative to the P/T junction. Specifically, the distance between the 5′ end of the primer strand and the “back face” of the loaded PCNA ring is decreased. In the initiation state for DNA synthesis, Pol δ directly engages 12 – 14 bp of the dsDNA region immediately upstream of the nascent P/T junction and PCNA encircles the next 10 – 12 bp directly upstream^24,32^. Hence, the aforementioned decrease in distance is likely due to the Pol δ•PCNA interaction maintaining loaded PCNA in closer proximity to the 5′ end of the primer strand. The results presented in **Figure 7D** – **F** indicate that during initiation of DNA synthesis by the resident Pol δ holoenzyme the interaction of RPA OBA is not observably affected and nearly all, if not all, RPA maintains contact(s) with a nascent P/T junction. In the initiation state for DNA synthesis, Pol δ directly engages the 3′ terminus of the primer strand and 4 - 6 nt of the template ssDNA strand immediately downstream of a nascent P/T junction and these protein•DNA interactions significantly overlap with those of the resident RPA complex (**Figure 1B**) ^12–18,24,32^. This situation is similar to that of the overlap of the activated loading complex and the RPA complex at nascent P/T junction where RPA OBD transiently releases from the 3′ terminus of the primer strand to accommodate binding and activation of the loading complex (**Figure 8**, *Step 1*). Thus, we postulate RPA OBD is released from the 3′ terminus of the primer strand to accommodate initiation of DNA synthesis by the assembled Pol δ holoenzyme. This hypothesis is currently being investigated.

Altogether, the results from the present study reveal that dynamic interactions of RPA subunits with a nascent P/T junction during assembly of a Pol δ holoenzyme and initiation of DNA synthesis both maintain RPA at a P/T junction and accommodate RFC, PCNA, and Pol δ. Furthermore, as discussed above, the only pathway for dissociation of loaded PCNA from a nascent P/T junction during these processes is through spontaneous opening of the PCNA ring, which is dramatically slow and essentially not observed. Hence, the coordinated actions of RPA, RFC, Pol δ and protein roadblocks maximize the efficiency of PCNA utilization throughout these processes. As Pol δ holoenzymes also carry out DNA synthesis during the major DNA repair and tolerance pathways, the current studies provide critical insights and direction for future studies on the DNA synthesis steps of long patch base excision repair, nucleotide excision repair, break-induced repair, mismatch DNA repair, translesion DNA synthesis and homology dependent recombination^2–6,11^.

## Methods

### Oligonucleotides

Oligonucleotides were synthesized by Integrated DNA Technologies (Coralville, IA) or Bio-Synthesis (Lewisville, TX) and purified on denaturing polyacrylamide gels. The concentrations of unlabeled DNAs were determined from the absorbance at 260 nm using the calculated extinction coefficients. The concentrations of Cy5-labeled DNAs were determined from the extinction coefficient at 650 nm for Cy5 (ε_650_ = 250,000 M^−1^cm^−1^). Concentrations of Cy3 labeled DNAs were determined from the extinction coefficient at 550 nm for Cy3 (ε_550_ = 136,000 M^−1^cm^−1^). For annealing two ssDNAs (as depicted in **Figure S1**), the primer and corresponding complementary template strands were mixed in equimolar amounts in 1X Annealing Buffer (10 mM TrisHCl, pH 8.0, 100 mM NaCl, 1 mM EDTA), heated to 95 °C for 5 minutes, and allowed to slowly cool to room temperature.

### Recombinant Human Proteins

Human RPA, Cy5-PCNA, exonuclease-deficient Pol δ (referred to herein as simply Pol δ) and RFC were obtained as previously described^21,47^. The concentration of active RPA was determined via a FRET-based activity assay as described previously^48^. Human RPA containing a Cy5 label at either residue 101 of the RPA2 subunit (Cy5-OBD-RPA) or residue 211 of the RPA1 subunit (RPA-OBA-Cy5) was obtained essentially as described for *S*. *cerevisiae* RPA^29^. Residue 101 of the RPA2 subunit and residue 211 of the RPA1 subunit reside in the OB-folds D and A of the RPA heterotrimeric complex, respectively. The concentration of active Cy5-labeled RPA was determined by a FRET-based assay that is described in detail in the **Supplementary Information**.

### Ensemble FRET Measurements

All experiments were performed at room temperature (23 ± 2 °C) and, unless indicated otherwise, in 1X Mg^2+^/Ca^2+^ buffer (20 mM HEPES, pH 7.5, 150 mM KCl, 5 mM MgCl_2_, 5 mM CaCl_2_) supplemented with 1 mM DTT, 1 mM ATP, and the ionic strength was adjusted to physiological (200 mM) by the addition of appropriate amounts of KCl. Ca^2+^ is included to account for experimental conditions in future studies and the presence of Ca^2+^ does not affect the amount of RPA that binds to ssDNA (**Figure S6**) nor the amount of PCNA loaded onto DNA by RFC (**Figure S4A** and **B** and **Table S1**). All experiments were performed in a 16.100F-Q-10/Z15 sub-micro fluorometer cell (Starna Cells) and monitored in a Horiba Scientific Duetta Bio fluorescence/absorbance spectrometer. Reaction solutions are excited at 514 nm and the fluorescence emission intensities (*I*) are monitored essentially simultaneously at 563 nm (*I*_563_) and 665 nm (*I*_665_) over time, recording *I* every 0.17 s. The acquisition rate of the instrument is 510,000 nm/min. Thus, for a given recording, the time between the acquisition of *I*_563_ and *I*_665_ for each time point is negligible (0.235 ms). For all FRET experiments, excitation and emission slit widths are 10 nm. For any recording of the fluorescence emission intensities (*I_665_* and *I_563_*), the approximate FRET efficiency is estimated from the equation 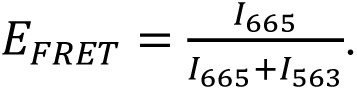 All recorded fluorescence emission intensities are corrected by a respective dilution factor and all time courses are adjusted for the time between the addition of each component and the fluorescence emission intensity recording (Δt ≤ 10 s). For each experiment below, the final concentrations of all reaction components are indicated. The concentrations of all nucleotides (ATP, ATPγS, dGTP) in all experimental solutions described below are each 1.0 mM and, hence, this concentration is maintained for each nucleotide upon mixing.

For PCNA loading experiments in the presence of ATPγS, a Cy3-labeled P/T DNA (20 nM, **Figure S1**), NeutrAvidin (80 nM), and ATPγS are pre-incubated with RPA (25 nM heterotrimer, wild type, RPA-OBA-Cy5 or Cy5-OBD-RPA) and the resultant solution is transferred to a fluorometer cell, and the cell is placed in the instrument. Fluorescence emission intensities (*I*_665_ and *I_563_*) are monitored over time until both signals stabilize for at least 1 min. Within this stable region, E_FRET_ values are calculated from the observed fluorescence emission intensities (*I*_665_ and *I_563_*) and averaged to obtain the E_FRET_ value observed prior to addition of the loading complex. Finally, a loading complex pre-formed with PCNA (20 nM homotrimer, Cy5-PCNA or PCNA), RFC (20 nM heteropentamer) and ATPγS is added, the resultant solution is mixed via pipetting, and the fluorescence emission intensities (*I*_665_ and *I_563_*) are monitored over time, beginning 10 s after the addition of loading complex (i.e., Δt ≤ 10 s).

For PCNA loading experiments in the presence of ATP, a Cy3-labeled P/T DNA (20 nM, **Figure S1**), NeutrAvidin (80 nM), and ATP are pre-incubated with RPA (25 nM heterotrimer, wild type, RPA-OBA-Cy5 or Cy5-OBD-RPA). Then, PCNA (20 nM homotrimer, Cy5-PCNA or PCNA) is added, the resultant solution is transferred to a fluorometer cell, and the cell is placed in the instrument. Fluorescence emission intensities (*I*_665_ and *I_563_*) are monitored over time until both signals stabilize for at least 1 min. Within this stable region, E_FRET_ values are calculated from the observed fluorescence emission intensities (*I*_665_ and *I_563_*) and averaged to obtain the E_FRET_ value observed prior to addition of the RFC•ATP. Finally, a pre-formed RFC•ATP complex (20 nM RFC heteropentamer) is added, the resultant solution is mixed via pipetting, and the fluorescence emission intensities (*I*_665_ and *I_563_*) are monitored over time, beginning 10 s after the addition of loading complex (i.e., Δt ≤ 10 s).

For Pol δ holoenzyme formation and initiation of DNA synthesis experiments with Cy5-PCNA, a solution containing a 5′ddPCy3/T DNA (20 nM, **Figure S1**), NeutrAvidin (80 nM), and ATP is pre-incubated with RPA (25 nM heterotrimer). Then, Cy5-PCNA (20 nM homotrimer) is added, the resultant solution is transferred to a fluorometer cell, and the cell is placed in the instrument. Fluorescence emission intensities (*I*_665_ and *I_563_*) are monitored over time until both signals stabilize for at least 1 min. Within this stable region, E_FRET_ values are calculated from the observed fluorescence emission intensities (*I*_665_ and *I_563_*) and averaged to obtain the E_FRET_ value observed prior to addition of the RFC•ATP. Next, a pre-formed RFC•ATP complex (20 nM RFC heteropentamer) is added, the resultant solution is mixed via pipetting, and the fluorescence emission intensities (*I*_665_ and *I_563_*) are monitored over time, beginning 10 s after the addition of loading complex (i.e., Δt ≤ 10 s). Fluorescence emission intensities (*I*_665_ and *I_563_*) are monitored over time until both signals stabilize for at least 1 min. Within this stable region, E_FRET_ values are calculated from the observed fluorescence emission intensities (*I*_665_ and *I_563_*) and averaged to obtain the E_FRET_ value observed prior to addition of Pol δ. Finally, Pol δ (20 – 80 nM Pol δ heterotetramer, ± dGTP) is added, the resultant solution is mixed by pipetting, and fluorescence emission intensities (*I*_665_ and *I_563_*) are monitored beginning ≤ 10 s after the addition of Pol δ (± dGTP).

For Pol δ holoenzyme formation and initiation of DNA synthesis experiments with PCNA, a solution containing ddP/5′TCy3 DNA (20 nM, **Figure S1**), NeutrAvidin (80 nM), and ATP is pre-incubated with RPA-OBA-Cy5 (25 nM heterotrimer). Then, PCNA (20 nM homotrimer) is added, followed by pre-formed RFC•ATP complex (20 nM RFC heteropentamer). The resultant solution is pre-incubated for the duration of RFC-catalyzed loading of PCNA (≥ 5 min), transferred to a fluorometer cell, and the cell is placed in the instrument. Fluorescence emission intensities (*I*_665_ and *I_563_*) are monitored over time until both signals stabilize for at least 1 min. Within this stable region, E_FRET_ values are calculated from the observed fluorescence emission intensities (*I*_665_ and *I_563_*) and averaged to obtain the E_FRET_ value observed prior to addition of the Pol δ. Next, Pol δ (40 nM Pol δ heterotetramer, ± dGTP) is added, the resultant solution is mixed via pipetting, and the fluorescence emission intensities (*I*_665_ and *I_563_*) are monitored over time, beginning 10 s after the addition of Pol δ (± dGTP, Δt ≤ 10 s). Fluorescence emission intensities (*I*_665_ and *I_563_*) are monitored over time until both signals stabilize for at least 1 min. Within this stable region, E_FRET_ values are calculated from the observed fluorescence emission intensities (*I*_665_ and *I_563_*) and averaged to obtain the E_FRET_ value observed after the addition of Pol δ. Finally, poly(dT)_70_ (413 nM) is added, the resultant solution is mixed by pipetting, and fluorescence emission intensities (*I*_665_ and *I_563_*) are monitored beginning ≤ 10 s after the addition of poly(dT)_70_.

## SUPPLEMENTAL INFORMATION

Supplemental Information includes Supplementary Results, Supplementary Methods, Supplemental Figures S1 – S14, and a Supplemental Table (Table S1).

## ACKNOWLEDGEMENTS

We would like to thank all members of the Hedglin and Antony labs for their efforts in reviewing/proofreading the current manuscript. This work was supported by funding from the National Institutes of Health to S.K. (F99CA274696), E.A. (R01 GM130756, R01 GM133967, R35 GM149320, and S10 OD030343) and M.H. (R35 GM147238-02),

## CONFLICTS OF INTEREST

The authors declare that they have no conflicts of interest with the contents of this article.^1^

## AUTHOR CONTRIBUTIONS

J.L.N., K.G.P., R.L.D., S.P., S.K., and V.K. expressed, purified, and characterized all proteins. J.L.N. and L.O.R. performed the experiments. M.H. designed the experiments. M.H. and E.A. analyzed the data and wrote the paper.

## Supplemental Information

### Supplementary Results

#### Experimental reaction buffer components do not affect the amount of RPA that binds to ssDNA nor the amount of PCNA loaded onto DNA by RFC

Calcium (Ca^2+^) is included in all experimental reaction buffers in the current study to account for experimental conditions in future studies. Rigorous control experiments were carried out to demonstrate that the results presented in the present study are not attributed to indirect effects of buffer components. We initially focused on RPA-DNA interactions. First, the concentration of active RPA heterotrimer was calculated as previously described in 1X Mg^2+^ buffer (25 mM HEPES, pH 7.5, 125 mM KOAc, 10 mM Mg(OAc)_2_) supplemented with 1 mM DTT, 1 mM ATP, and ionic strength adjusted to 200 mM by addition of KOAc^1^. The ssDNA substrate utilized in these assays (poly(dT)_30_-FRET) accommodates one RPA heterotrimer and is shown in **Figure S1**. Next, these titrations were repeated in the same experimental reaction buffer except with amended amounts of RPA heterotrimer that accounted for the calculated concentration of active RPA heterotrimer (**Figure S2A**) and the data (E_FRET_ values) is plotted as a function of the ratios of the concentrations of active RPA heterotrimer and DNA (i.e., [RPA]:[DNA], **Figure S2B**). As expected, saturation occurs at a ratio of 1:1 (1.017 ± 0.0545 [RPA]:[DNA]). Finally, these experiments were repeated with the same preparation of human RPA heterotrimer except in 1X Mg^2+^/Ca^2+^ buffer. In this Mg^2+^/Ca^2+^ buffer, Ca^2+^ is included in addition to magnesium (Mg^2+^) and the acetate (OAc^-^) from Mg^2+^ buffer is replaced with chloride (Cl^-^). As observed in **Figure S2C**, saturation occurs in 1X Mg^2+^/Ca^2+^ buffer at ratio (0.938 ± 0.0507 [RPA]:[DNA]) that is within experimental error of that observed in 1X Mg^2+^ buffer (1.017 ± 0.0545 [RPA]:[DNA], **Figure S2C**). This indicates that experimental reaction buffer components do not affect the amount of RPA that binds to ssDNA. Next, we focused on RFC-catalyzed loading of PCNA onto a P/T junction.

RFC-catalyzed loading of PCNA onto nascent P/T junctions that are engaged with RPA was monitored exactly as described in **Figure 3** in the main text. Under the conditions of the assay, PCNA loading is biphasic and stoichiometric^2,3^. Shown in **Figure S3A** – **B** is the data observed in 1X Mg^2+^ buffer. Upon addition of RFC, the observed changes in *I*_665_ and *I*_563_ are synchronized and anti-correlated (**Figure S3A**, *Top*), indicating the appearance and increase in FRET (**Figure S3A**, *Bottom*). As observed in **Figure S3B**, E_FRET_ traces rapidly increase to values significantly above the E_FRET_ values observed for no interaction between Cy5-PCNA and the 5′ddPCy3/T•RPA complex. As expected, the rapid increase in E_FRET_ observed upon addition of RFC•ATP is biphasic with observed rate constants of *k*_obs inc,1_ = 0.122 ± 0.004 s^-1^ and *k*_obs inc,2_ = 1.28 ± 0.05 (x 10^-2^) s^-1^ (reported in **Table S1**) and an overall (i.e., total) amplitude of A_T_ = 0.351 ± 0.019. Importantly, the rate constants observed in **Figure S3B** agree very well with values reported in a previous study (*k*_obs inc,1_ = 0.134 ± 0.010 s^-1^ and *k*_obs inc,2_ = 2.03 ± 0.03 (x 10^-2^) s^-1^) that analyzed PCNA loading under similar conditions by monitoring sensitized Cy5 acceptor fluorescence emission intensity (*I*_665_) via stopped flow. Under these conditions, *k*_obs inc_,_1_ describes a kinetic step along the PCNA loading pathway that occurs prior to and much slower than binding of the loading complex to the nascent P/T junction. *k*_obs inc_,_2_ describes release of the RFC•ADP complex into solution via dissociation from RPA engaged with the nascent P/T junction^2^. The overall amplitude (A_T_) indicates the overall increase in FRET observed upon stable loading of a single Cy5-PCNA onto each 5′ddPCy3/T DNA substrate. Shown in **Figure S3C** – **D** is the data observed in 1X Mg^2+^/Ca^2+^ buffer. E_FRET_ traces observed upon addition of RFC•ATP are also biphasic. Importantly, the overall amplitude observed in **Figure S3D** (A_T_ = 0.368 ± 0.014) is within experimental error of that observed in the alternative buffer (A_T_ = 0.351 ± 0.019, **Figure S3B**), indicating that experimental reaction buffer components do not affect the amount of PCNA loaded onto DNA by RFC. *k*_obs_,_2_ observed in **Figure S3D** (*k*_obs inc, 2_ = 1.72 ± 0.02 (x 10^-2^) s^-1^, **Table S1**) agrees very well with that observed in **Figure S3B** for the alternative buffer (*k*_obs inc_,_2_ = 2.03 ± 0.03 (x 10^-2^) s^-1^, **Table S1**) suggesting that *k*_obs inc_,_2_ under these conditions also represents release of the RFC•ADP complex into solution via dissociation from RPA engaged with the nascent P/T junction. *k*_obs inc_,_1_ observed for each experimental reaction buffer are within 2-fold of each other. This suggests either that *k*_obs inc,1_ describes either; 1) the same kinetic step(s) for each condition and, hence, this step is dependent on the experimental reaction buffer components; or 2) a distinct step(s) for each condition. Regardless, the identical total amplitudes (A_T_) observed in **Figure S3** indicate that both experimental reaction buffers fully support RFC-catalyzed loading of PCNA onto P/T junctions.

#### RPA rapidly engages a nascent P/T junction in a stable interaction

To investigate RPA/DNA interactions at P/T junctions, we designed and carried out FRET assays utilizing Cy3, Cy5 FRET pairs comprised of a Cy3-labeled P/T DNA substrate (Figure S1) and a corresponding Cy5-labeled RPA. To monitor interactions of RPA OBA with P/T DNA we utilized a substrate (Figure S4A, ddP/5′TCy3 DNA) with a 3′ dideoxy-terminated primer and a Cy3 donor near the 5′ end of the template strand and RPA with a Cy5 acceptor on OBA (Figure S4B, RPA-OBA-Cy5) that faces the Cy3 donor (Figure S4C)^4–11^. To monitor interactions of RPA OBD with P/T DNA we utilized a substrate (Figure S4D, 3′PCy3/T) with a Cy3 donor on the 3′ terminus of the primer strand and RPA with a Cy5 acceptor on OBD (Figure S4E, Cy5-OBD-RPA) that faces the Cy3 donor (Figure S4F)^12^. Each Cy3-labeled P/T junction was tested for stable interaction with the corresponding Cy5-labeled RPA by monitoring the FRET signals observed at equilibrium after excitation with 514 nm light. Here, Cy5 on RPA can be excited via FRET from Cy3 on a P/T junction only when the two cyanine labels remain in close proximity of each other (i.e., ≤ 10 nm), as depicted in Figures S4C and S4F. This is indicated by an increase in the fluorescence emission intensity of the Cy5 acceptor at 665 nm (Cy5 acceptor fluorescence emission maximum, *I*_665_) and a concomitant decrease in the fluorescence emission intensity of the Cy3 donor at 563 nm (Cy3 donor fluorescence emission maximum, *I*_563_). A FRET signal is clearly observed for both Cy3, Cy5 FRET pairs only when both the respective Cy3-labeled P/T junction and the corresponding Cy5-labeled RPA are present (Figure S4G and S4H). Collectively, this indicates that RPA engages a P/T junction in a stable interaction. This equilibrium FRET assay was adapted to determine the concentrations of active Cy5-labeled RPAs (Figure S5).

To analyze the kinetics of RPA binding to nascent P/T junctions (Figure S6A), *I_563_* and *I*_665_ are first monitored over time for ddP/5’TCy3 DNA alone. Then, the fluorescence emission intensity recording is paused, RPA-OBA-Cy5 is added to the reaction mixture, the resultant solution is mixed, and the fluorescence emission intensity recording is resumed within 10 s of the addition (i.e., “dead time” ≤ 10 s). Upon addition of RPA-OBA-Cy5, *I*_665_ rapidly increases concomitantly with a rapid decrease in *I*_563_ (Figure S6B, *Top*) after which both fluorescence emission intensities stabilize and persist over time. These synchronized, anti-correlated changes in *I*_563_ and *I*_665_ are indicative of the appearance and increase in FRET (Figure S6B, *Bottom*). As observed in Figure S6C, E_FRET_ traces rapidly increase to values significantly above the E_FRET_ traces predicted for no interaction between RPA-OBA-Cy5 and ddP/5′TCy3 DNA. Furthermore, the rapid increase in E_FRET_ observed upon addition of RPA-OBA-Cy5 is comprised of at least two phases (i.e., biphasic) with an observed rate constant for the slower phase (*k*_obs inc_,_2_) of 9.45 x 10^-3^ + 2.23 x 10^-3^ s^-1^. Similar results are observed for the 3′PCy3/T DNA, Cy5-OBD-RPA FRET pair (Figure S6D – F) except that association of the Cy5-labeled OBD of RPA with the Cy3-labeled P/T junction is completed within the dead time of the experiment (≤ 10 s). This behavior agrees with previous studies that observed differential kinetics for binding of OBA and OBD to ssDNA in the context of the complete, heterotrimeric RPA complex^12–14^. Altogether, the results presented in Figures S4 and S6 indicate that RPA rapidly engages P/T junctions in a stable interaction.

#### PCNA is loaded onto a P/T junction in 1:1 stoichiometry regardless of the position of the Cy3 label

In order to investigate the effects of the Cy3 donor located at the 3′ primer terminus of the of the 3′PCy3/T DNA substrate (**Figure S1**) on RFC-catalyzed loading of PCNA, we carried out PCNA loading experiments under two different conditions. First, we investigated the kinetics of PCNA loading by carrying out PCNA loading reactions in the presence of ATP exactly as described in in the main text except with 3′Cy3P/T DNA (**Figure S9A**). In short, 20 nM 3′PCy3/T DNA (pre-bound with 80 nM NeutrAvidin and 1 mM ATP are pre-incubated with 25 nM wild type RPA. Then, 20 nM Cy5-PCNA is added. PCNA loading is initiated by the addition of 20 nM pre-formed RFC•ATP complex and monitored via FRET over time (**Figure S9B**). Under these conditions the concentrations of RFC, PCNA, and P/T DNA are stoichiometric (i.e., 1:1:1). As observed in **Figure S9C**, E_FRET_ traces rapidly increase to values significantly above the E_FRET_ values observed for no interaction between Cy5-PCNA and the 3′PCy3/T•RPA complex. The rapid increase in E_FRET_ observed upon addition of RFC is biphasic with observed rate constants of *k*_obs inc,1_ = 4.42 ± 0.10 (x 10^-2^) s^-1^ and *k*_obs inc,2_ = 1.59 ± 0.09 (x 10^-2^) s^-1^ (**Table S1**). The rate constants observed for the 3′Cy3P/T DNA substrate (**Figure S9C**, **Table S1)** are nearly identical to those observed for the 5′Cy3P/T DNA substrate (**Table S1**), indicating that the Cy3 label on the 3′ terminus of the primer of the 3′Cy3P/T DNA substrate does not affect kinetics of RFC-catalyzed loading of PCNA onto P/T junctions. The overall total amplitude (A_T_) observed for the 3′Cy3P/T DNA substrate (**Figure S9C**) is > 2-fold less than that observed for the 5′Cy3P/T DNA substrate, likely due to the Cy5 label on PCNA being oriented away from the Cy3 donor (rather than towards) when Cy5-PCNA is loaded onto the 3′Cy3P/T DNA substrate by RFC. Next, we investigated the stoichiometry of PCNA loading by characterizing the overall total amplitudes (A_T_) of PCNA loading via a titration of the E_FRET_ signal observed at equilibrium (i.e., when A_T_ has been achieved). In short, a Cy3-labeled P/T DNA (55 nM of either 5′Cy3P/T or 3′Cy3P/T pre-bound with 220 nM NeutrAvidin) is pre-saturated with wild type RPA prior to the addition of RFC (55 nM). Cy5-PCNA is then titrated in and the equilibrium E_FRET_ is monitored. As expected for the 5′PCy3/T DNA substrate, E_FRET_ increased linearly with Cy5 PCNA up to a Cy5-PCNA:Cy3-DNA ratio of 1:1 after which the E_FRET_ values remain constant (**Figure S10**)^2,3^. This confirms the validity of the approach and that RFC-catalyzed loading of PCNA onto the 5′PCy3/T DNA substrate is stoichiometric. For the 3′PCy3/T DNA substrate, identical behavior is observed. This indicates that the location of the Cy3 label on the primer strand does not affect the amount of PCNA loaded onto DNA by RFC. Together, the results presented in **Figures S9** and **S10** demonstrate that the Cy3 donor located at the 3′ primer terminus of the of the 3′PCy3/T DNA substrate has no effect on RFC-catalyzed loading of PCNA on the resident P/T junction.

#### dGTP inhibits RFC-catalyzed re-loading of PCNA onto a P/T junction

In the current study, RFC-catalyzed loading of PCNA onto a P/T junction in the presence of ATP is stoichiometric. After dissociation of the RFC•ADP complex into solution, the loaded PCNA is left behind on the duplex region of the P/T junction and randomly and rapidly diffuses along the double-stranded DNA (dsDNA)^15^. Recent *in vivo* evidence suggests that enzyme catalyzed unloading of PCNA from a P/T junction will not occur until the primer is completely extended and ligated to the downstream duplex region^16^. Physical blocks i.e., “protein roadblocks,” restrict translocation of PCNA away from the P/T junction via diffusion. Diffusion of loaded PCNA along the duplex region is restricted by high-affinity DNA-binding proteins, such as histones and transcription factors, that rapidly bind nascent dsDNA upstream of P/T junctions during S-phase of the cell cycle when DNA replication occurs. This is emulated in the current study by the biotin-NeutrAvidin complex at the blunt duplex end of the DNA substrates. RPA engaged with the ssDNA downstream of nascent P/T junctions prohibits diffusion of loaded PCNA along ssDNA as well as RFC-catalyzed unloading of PCNA^2,3^. In the absence of catalyzed unloading of PCNA and significant translocation of loaded PCNA, the only pathway for dissociation of loaded PCNA from a nascent P/T junction is through spontaneous opening of the PCNA ring, which is dramatically slow [*k*_open_ = 1.25 ± 0.32 (x 10^-3^) s^−1^]^17^, ∽20-fold slower than the observed rate constant RFC-catalyzed loading of “free” PCNA onto a P/T junction in the presence of ATP (**Figure 3, 5, S3, S3 – S9**, and **Table 1**). This suggests that, upon dissociation of PCNA from a nascent P/T junction via spontaneous opening of the PCNA ring, RFC utilizes ATP to instantly reload PCNA back onto the nascent P/T junction such that the loss of loaded PCNA from a nascent P/T junction and, hence, E_FRET_, is not observed^2,18^. To directly confirm this, we continuously analyzed DNA-PCNA interactions over time as reaction conditions progressively evolved (**Figure S11A**). First, the 5′ddPCy3/T DNA substrate is pre saturated with native RPA, Cy5-PCNA is added, and *I_563_* and *I*_665_ are monitored over time. Next, pre-formed RFC•complex is added to the and the fluorescence emission intensities are monitored over time until PCNA loading is complete. Under the conditions of the assay, PCNA loading is stoichiometric and biphasic^2^. Finally, unlabeled PCNA is added in excess and the fluorescence emission intensities are monitored over time. Under these conditions, once loaded Cy5-PCNA dissociates from the Cy3-labeled P/T junction via spontaneous opening of the PCNA ring, re-loading of “free” Cy5-PCNA is prohibited due to the 70-fold excess of unlabeled PCNA, and consequently the observed FRET decreases. Here, the disappearance of FRET is rate-limited by and directly reports on spontaneous opening of the PCNA ring.

Upon addition of RFC•ATP, *I*_665_ rapidly increases concomitantly with a rapid decrease in *I*_563_ after which both fluorescence emission intensities stabilize and persist over time (Figure S11B, *Top*). These synchronized, anti-correlated changes in *I*_563_ and *I*_665_ are indicative of the appearance and increase in FRET (Figure S11B, *Bottom*). As expected, E_FRET_ traces observed in Figure S11C rapidly increase in a biphasic manner upon addition of RFC•ATP and plateau at values significantly above the E_FRET_ traces observed for no interaction between Cy5-PCNA and 5′ddPCy3/T. At this point, a Cy5-PCNA has been loaded onto each 5′ddPCy3/T DNA and RFC•ADP has released into solution and exchanged ADP for ATP. Both observed rate constants for RFC-catalyzed loading of PCNA in the presence of ATP are in excellent agreement with results from the current study obtained under the same experimental conditions (Table S1). *k*_obs inc, 1_ (in Figure S11C), which encompasses all kinetic steps along the PCNA loading pathway up to and including release of loaded PCNA onto P/T DNA, is 4.46 + 0.29 (x 10^-2^) s^-1^. *k*_obs inc,2_ (in Figure S11C), which reports on release of RFC•ADP into solution (via its dissociation from the resident RPA engaged with the P/T junction) is 1.70 + 0.12 (x 10^-2^) s^-1^.

Upon addition of excess, unlabeled PCNA, the observed changes in *I*_665_ and *I*_563_ are synchronized and anti-correlated (**Figure S11B**, *Top*), indicating a decrease in FRET (**Figure S11B**, *Bottom*). As observed in **Figure S11C**, upon addition of excess unlabeled PCNA, E_FRET_ traces decrease to E_FRET_ values observed for no interaction between Cy5-PCNA and 5′ddPCy3/T DNA (i.e., all Cy5-PCNA remaining “free” in solution) indicating that all Cy5-PCNA dissociates from the 5′ddPCy3/T•RPA complex. Furthermore, the decrease in E_FRET_ observed upon addition of excess unlabeled PCNA is comprised of a single phase (i.e., monophasic) with an observed rate constant [*k*_obs dec_ = 1.93 ± 0.01 (x 10^-3^) s^-1^] that is in excellent agreement with the rate constant for spontaneous opening of the PCNA ring [*k*_open_ = 1.25 ± 0.32 (x 10^-3^) s^−1^]^17^ and is 23.1 ± 1.50-fold slower than the rate constant for RFC-catalyzed loading of “free” PCNA onto a P/T junction in the presence of ATP [*k*_obs inc, 1_ = 4.46 ± 0.29 (x 10^-2^) s^-1^]. Altogether, this confirms that; 1) dissociation of PCNA from nascent P/T junctions is governed entirely by spontaneous opening of the PCNA ring; and 2) upon dissociation of PCNA from a nascent P/T junction, RFC•ATP instantly reloads PCNA back onto the nascent P/T junction such that the loss of loaded PCNA from a nascent P/T junction is not observed. In other words, RFC together with ATP continuously maintain loaded PCNA on all P/T junctions.

Interestingly, when the experiments described in Figure S11 and Figure 6 (in the main text) were repeated by replacing unlabeled PCNA and Pol δ, respectively with dGTP (1 mM final concentration, Figure S12A), *I*_665_ decreases slightly over time concomitantly with a slight increase in *I*_563_ over time (Figure S12B, *Top*). These synchronized, anti-correlated changes in *I*_563_ and *I*_665_ are indicative of a decrease in FRET (Figure S12B, *Bottom*). As observed in Figure S12C, upon addition of dGTP, E_FRET_ traces very slightly, but reproducibly, decrease (% FRET Change = -11.7 + 3.7 %) with an observed rate constant [*k*_obs dec_ = 3.75 + 0.41 (x 10^-3^) s^-1^] that agrees with the rate constant observed in Figure S11C [*k*_obs dec_ = 1.93 + 0.01 (x 10^-3^) s^-1^] for PCNA unloading via spontaneous opening of the PCNA ring. This suggests that; 1) PCNA dissociates from the P/T junction under these conditions via spontaneous opening of the PCNA ring; and 2) dGTP (at a concentration of 1 mM) inhibits RFC-catalyzed reloading of PCNA back onto the P/T junction. Regarding the latter point, previous studies of human RFC observed that dGTP decreases DNA-dependent nucleotide triphosphate hydrolysis by RFC approximately 4 fold and significantly tempers PCNA-dependent stimulation of Pol δ-mediated DNA synthesis by ∽8-fold, suggesting that dGTP significantly inhibits loading of PCNA onto P/T junctions by RFC^19^. To directly test this, we repeated PCNA loading assays (described in Figures 3, 5, S3, and S7 – S9) in the presence of dGTP (Figure S13).

Upon addition of RFC•dGTP, the observed changes in *I*_665_ and *I*_563_ are synchronized and correlated (Figure 13B, *Top*). This behavior is due to nonspecific effects^20^. For the example FRET trajectory depicted in Figure 13B (*Bottom*), E_FRET_ values observed prior to the addition of the RFC•dGTP complex are maintained after addition of the RFC•dGTP loading complex. For the averaged E_FRET_ trajectory depicted in Figure S13C, the E_FRET_ traces observed prior to the addition of the RFC•dGTP complex persist and are maintained at the E_FRET_ values observed for no interaction between Cy5-PCNA and the 5′ddPCy3/T•RPA complex. This indicates that dGTP (at a concentration of 1 mM) prohibits loading of PCNA onto P/T junctions by RFC under these conditions and, hence, likely inhibits re-loading of PCNA when it is added (at a concentration of 1 mM) after a significant incubation time (in Figure S12 and Figure 6) where ATP has been depleted and ADP has increased through spontaneous hydrolysis of ATP and RFC-catalyzed PCNA loading and re-loading.

#### Human RPA undergoes facilitated exchange with free ssDNA

The human RPA complex has exceptionally high affinity for ssDNA at physiological ionic strength but can undergo facilitated exchange due to the dynamic ssDNA-binding interactions of its individual OB folds, enabling the human RPA complex to rapidly exchange between free and ssDNA-bound states when free, high-affinity ssDNA-binding proteins are present in solution. In this process, human RPA complexes exist in microscopically dissociated states that only undergo macroscopic dissociation when free, high-affinity ssDNA-binding proteins are available to occupy the ssDNA that is exposed during the microscopic dissociation events^12,13,21,22^. Accordingly, this concentration-dependent RPA turnover should also be observed when free ssDNA is present in solution. Here, microscopically dissociated states of RPA complexes only undergo macroscopic dissociation when free ssDNA sequences are available to occupy the OB folds of RPA that are exposed during the microscopic dissociation events. This behavior is akin to intersegmental transfer of DNA-binding proteins that contain at least two DNA binding domains^23^. To test this, we analyzed the kinetics of RPA dissociation from nascent P/T junctions (**Figure S14A**).

First, ddP/5’TCy3 DNA is pre-incubated with RPA-OBA-Cy5 at a ratio of 1:1 and *I_563_* and *I*_665_ of the resultant mixture are monitored over time. Then, the fluorescence emission intensity recording is paused, buffer containing varying concentrations of free poly(dT)_70_ is added to the reaction mixture, the resultant solution is mixed, and the fluorescence emission intensity recording is resumed within 10 s of the addition (i.e., “dead time” ≤ 10 s). The ssDNA binding affinity of human RPA is highest for poly(dT) and each poly(dT)_70_ accommodates at least two RPA complexes^4–6,24^. Hence, poly(dT)_70_ serves as an effective trap to release RPA-OBA-Cy5 from the ddP/5′TCy3 DNA substrate via facilitated exchange and prohibit re-binding. Upon addition of 0.25 μM poly(dT)_70_, the observed changes in *I*_665_ and *I*_563_ are synchronized and anti correlated (**Figure S14B**, *Top*), indicating a decrease in FRET (**Figure S14B**, *Bottom*). As observed in **Figure S14C**, E_FRET_ traces observed in the presence of 0.25 μM poly(dT)_70_ rapidly decrease to the E_FRET_ traces predicted for no interaction between RPA-OBA-Cy5 and ddP/5′TCy3 DNA. Furthermore, the rapid decrease in E_FRET_ observed upon addition of poly(dT)_70_ is comprised of at least two phases (i.e., biphasic) with an observed rate constants of *k*_obs dec_,_1_ = 9.02 + 0.49 (x10^-2^) s^-1^ and *k*_obs dec_,_2_ = 7.09 + 0.13 (x10^-3^) s^-1^. As expected, RPA-OBA-Cy5 remained tightly bound to the ddP/5’TCy3 DNA in mock reactions lacking poly(dT)_70_ (**Figure S14D**). In contrast, RPA-OBA-Cy5 rapidly and completely dissociated from the ddP/5’TCy3 DNA at each concentration of poly(dT)_70_ (**Figure S14D**) and the rates RPA-OBA-Cy5 dissociation are dependent upon the concentration of poly(dT)_70_ (**Figure S14E**), indicating a second order, biomolecular reaction. Altogether, this indicates that human RPA complexes can undergo concentration-dependent facilitated exchange between ssDNA sequences.

### Supplementary Methods

#### Determining stoichiometry of RPA:ssDNA interactions via FRET

The Cy3/Cy5-labeled ssDNA oligonucleotide (poly(dT)_30_-FRET) accommodates one RPA heterotrimer and is shown in **Figure S1**^25^. All experiments were performed at room temperature (23 ±2 °C) in either 1X Mg^2+^/Ca^2+^ buffer and the ionic strength adjusted to 200 mM by addition of KCl or in 1X Mg^2+^ buffer with the ionic strength adjusted to 200 mM by addition of KOAc. The excitation and emission slit widths were set to 5 nm. The poly(dT)30-FRET ssDNA is titrated with increasing concentrations of RPA. For each RPA addition, fluorescence emission intensities (*I*_665_ and *I_563_*) are monitored over time until both signals stabilize for at least 1 min. Within this stable region, E_FRET_ values are calculated from the observed fluorescence emission intensities (*I*_665_ and *I_563_*) and averaged to obtain the final E_FRET_ value for a given RPA addition. Under the experimental conditions, RPA binding is stoichiometric and, hence, E_FRET_ increases linearly until the ssDNA is saturated with Cy5-labeled RPA (i.e., equivalence point)^12,25^. Data is plotted as a function of the ratio of the concentrations of active RPA and DNA (i.e., [RPA]:[DNA]) and fit to two segment lines (a linear regression with a positive slope and a flat line). The equivalence point (RPA per DNA) is calculated from the intersection of the two segment lines.

#### Equilibrium FRET assays to characterize RPA interactions

One or more of the following components are pre-equilibrated in a fluorometer cell with 1 mM ATP; Cy3-labeled P/T DNA (20 nM, **Figure S1**), NeutrAvidin (80 nM), and a Cy5-labeled RPA (25 nM heterotrimer, RPA-OBA-Cy5 or Cy5-OBD-RPA). The cell is subsequently placed in the instrument, the respective solution is excited at 514 nm, and fluorescence emission spectra (530 nm – 750 nm) are recorded (1 spectra/25.9 ms) until the spectra stabilizes for at least 1 min. Spectra observed within this stable period are averaged to obtain the final spectra for the respective condition.

#### Equilibrium FRET assays to determine the concentration of active human Cy5-labeled RPA

A Cy3-labeled Bio-P/T DNA substrate (**Figure S1**) is titrated with increasing concentrations of a human Cy5-labeled RPA. For each RPA addition, fluorescence emission intensities (*I*_665_ and *I_563_*) are monitored over time until both signals stabilize for at least 1 min. Within this stable region, E_FRET_ values are calculated from the observed fluorescence emission intensities (*I*665 and *I_563_*) and averaged to obtain the final E_FRET_ value for a given RPA addition. Under the experimental conditions, RPA binding is stoichiometric and, hence, E_FRET_ increases linearly until the ssDNA is saturated with Cy5-labeled RPA (i.e., equivalence point)^12,25^. Data is fit to two segment lines (a linear regression with a positive slope and a flat line) and the equivalence point is calculated from the intersection of the two segment lines. The concentration of the active Cy5-labeled RPA (in μM) is determined by dividing the amount of RPA-binding sites (in pmoles) by the total volume (in μL) of Cy5-labeled RPA at the equivalence point. Each Cy3-labeled P/T DNA substrate utilized in the present study accommodates 1 RPA heterotrimer. Hence, for a given Cy3-labeled P/T DNA, Cy5-labeleld RPA FRET pair, the amounts of RPA-binding sites is equal to the amount of the respective Cy3-labeled P/T DNA.

#### Pre-steady state FRET assays to monitor RPA-DNA interactions

A solution containing a Cy3-labeled P/T DNA (20 nM, **Figure S1**), NeutrAvidin (80 nM, Thermo Scientific), and ATP (1 mM, Thermo Scientific) is pre-incubated, and the resultant solution is transferred to a fluorometer cell that is then placed in the instrument. Fluorescence emission intensities (*I*_665_ and *I_563_*) are monitored over time until both signals stabilize for at least 1 min. Within this stable region, E_FRET_ values are calculated from the observed fluorescence emission intensities (*I*_665_ and *I_563_*) and averaged to obtain the E_FRET_ value observed prior to addition of Cy5-labeled RPA. Then, a Cy5-labeled RPA (25 nM heterotrimer, either Cy5-OBD-RPA or RPA-OBA-Cy5) is added, the resultant solution is mixed by pipetting, and the fluorescence emission intensities (*I*_665_ and *I_563_*) are monitored over time, beginning 10 s after the addition of the Cy5-labeled RPA (i.e.,Δt <u>≤</u> 10 s). To determine the predicted E_FRET_ trace for a Cy5-labeled RPA remaining completely disengaged from a Cy3-labeled P/T DNA, these experiments are repeated with Cy5-labeled RPA alone and with a Cy3-labeled P/T DNA alone and the E_FRET_ is calculated for each time point by the equation

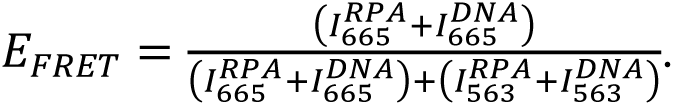

#### Equilibrium FRET assays to determine the stoichiometry of RFC-catalyzed loading of PCNA onto P/T junctions

A Cy3-labeled P/T DNA (55 nM of either 5′Cy3P/T or 3′Cy3P/T pre-bound with 220 nM NeutrAvidin) is saturated with RPA prior to the addition of RFC (55 nM) and then Cy5-PCNA is titrated in. After each addition of Cy5-PCNA, E_FRET_ is monitored over time until the signal stabilizes for at least 1 min. E_FRET_ values observed within this stable period are averaged to obtain the E_FRET_ value observed for the respective addition of Cy5-PCNA.

*Pre-steady state FRET assays to monitor PCNA unloading*. A solution containing a 5′ddPCy3/T DNA (20 nM, **Figure S1**), NeutrAvidin (80 nM), and ATP (1 mM) is pre-incubated with RPA (25 nM heterotrimer). Then, Cy5-PCNA (20 nM homotrimer) is added, the resultant solution is transferred to a fluorometer cell, and the cell is placed in the instrument. E_FRET_ is monitored over time until the signal stabilizes for at least 1 min. E_FRET_ values observed within this stable period are averaged to obtain the E_FRET_ value observed prior to addition of RFC•ATP. Next, a pre formed RFC•ATP complex (20 nM RFC heteropentamer, 1 mM ATP) is added, the resultant solution is mixed via pipetting, and E_FRET_ is monitored beginning <u>≤</u> 10 s after the addition of RFC (i.e., Δt <u>≤</u> 10 s) and continues until the signal stabilizes for at least 1 min. E_FRET_ values observed within this stable period are averaged to obtain the E_FRET_ value observed prior to addition of unlabeled PCNA. Finally, unlabeled PCNA (1.4 μM homotrimer) is added, the resultant solution is mixed by pipetting, and E_FRET_ is monitored beginning <u>≤</u> 10 s after the addition of unlabeled PCNA (Δt <u>≤</u> 10 s).

#### Pre-steady state FRET assays to monitor facilitated exchange of RPA between ssDNA sequences

A solution containing ddP/5′TCy3 (50 nM, **Figure S1**), NeutrAvidin (200 nM, Thermo Scientific), and ATP (1 mM, Thermo Scientific), and RPA-OBA-Cy5 (55 nM heterotrimer) is pre-incubated, and the resultant solution is transferred to a fluorometer cell that is then placed in the instrument. Fluorescence emission intensities (*I*_665_ and *I_563_*) are monitored over time until both signals stabilize for at least 1 min. Within this stable region, E_FRET_ values are calculated from the observed fluorescence emission intensities (*I*_665_ and *I_563_*) and averaged to obtain the E_FRET_ value observed prior to addition of poly(dT)_70_ . Then, poly(dT)_70_ (0.0 – 2.25 μM) is added, the resultant solution is mixed by pipetting, and the fluorescence emission intensities (*I*_665_ and *I_563_*) are monitored over time, beginning 10 s after the addition of the poly (dT)_70_ (i.e., Δt <u>≤</u> 10 s).

### Supplemental Figures

**Figure S1.**
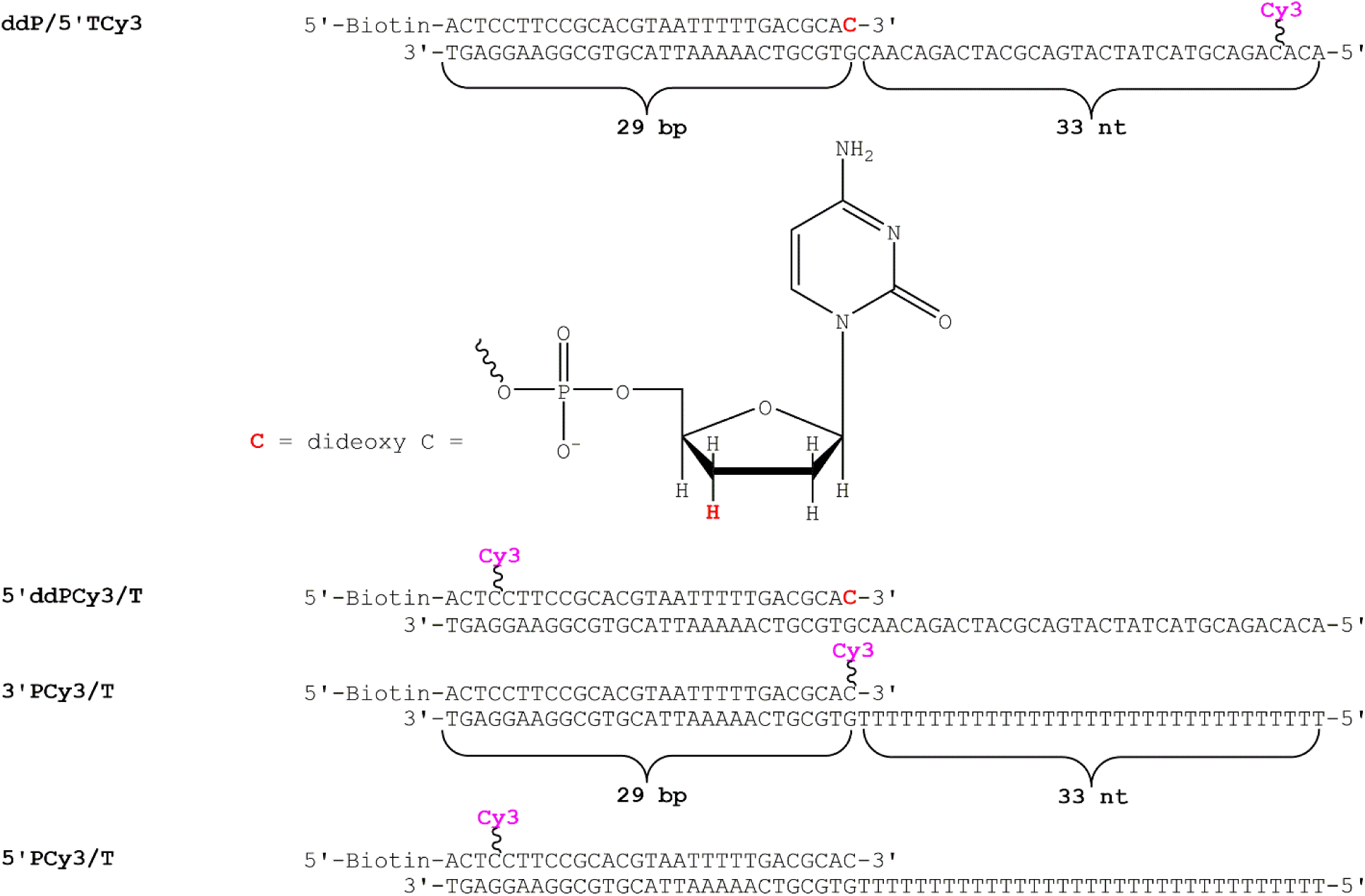
DNA substrates utilized in this study. For substrates containing dsDNA regions, the sequences and lengths (29 bp) of these regions are all identical. When annealed, each substrate mimics a nascent P/T junction. The size of the dsDNA P/T region (29 bp) is in agreement with the requirements for assembly of a PCNA ring onto DNA by RFC^2,3,18^. The ssDNA regions adjacent to the 3′ end of the P/T junctions are 33 nt in length and accommodate one RPA heterotrimer^4–6^. RPA prevents loaded PCNA from sliding off the ssDNA end of the substrate^2^. When pre-bound to NeutrAvidin, the biotin attached to the 5′-end of a primer strand prevents loaded PCNA from sliding off the dsDNA end of the substrate. ssDNA comprised only of T (i.e., poly(dT)_X_) is incapable of adapting stable secondary structures^26^. Primers terminated at the 3′ end with a dideoxy C nucleotide cannot be extended by Pol δ.

**Figure S2.**
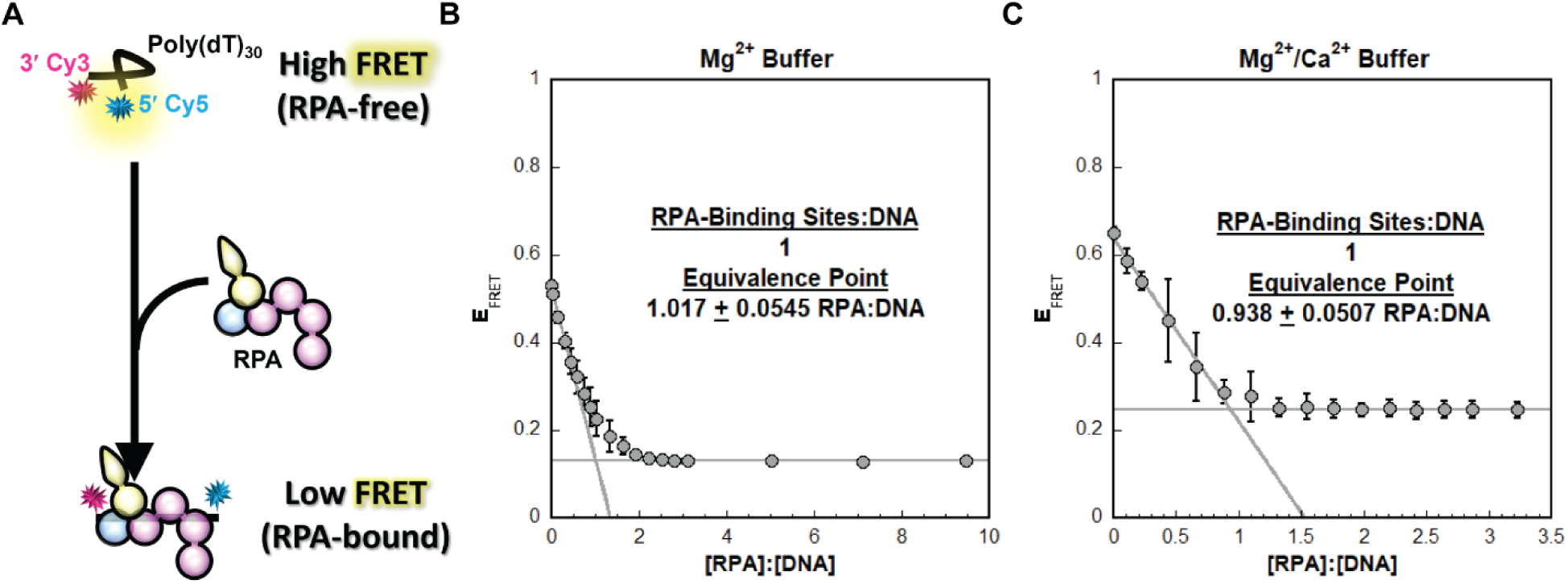
Effect of experimental reaction buffers on RPA binding ssDNA. (**A**) Schematic representation of the FRET experiment utilizing poly(dT)_30_ FRET ssDNA and native RPA. Poly(dT)_30_ FRET is terminally labeled with a 3′ Cy3 (FRET donor) and a 5′ Cy5 (FRET acceptor). In the absence of RPA, free DNA (poly(dT)_30_-FRET) forms a compact, flexible structure, bringing the two cyanine fluorophores close together and yielding a high E_FRET_. Binding of an RPA stretches the engaged ssDNA and increases its bending 2 – 3 fold^12,27^, thereby increasing the Cy3, Cy5 distance and reducing E_FRET_. (**B** - **C**) FRET data for RPA titrations carried out in Mg^2+^ Buffer (panel **B**) and Mg^2+^/Ca^2+^ Buffer (panel **C**). For each, poly(dT)_30_-FRET (10 nM) is titrated with RPA and E_FRET_ is monitored. The observed E_FRET_ is plotted as a function of the ratio of concentrations of active RPA and DNA (i.e., [RPA]:[DNA]). Each data point represents the mean ± S.E.M. of at least three independent measurements. The equivalence points for each panel are indicated with standard errors of the calculations.

**Figure S3.**
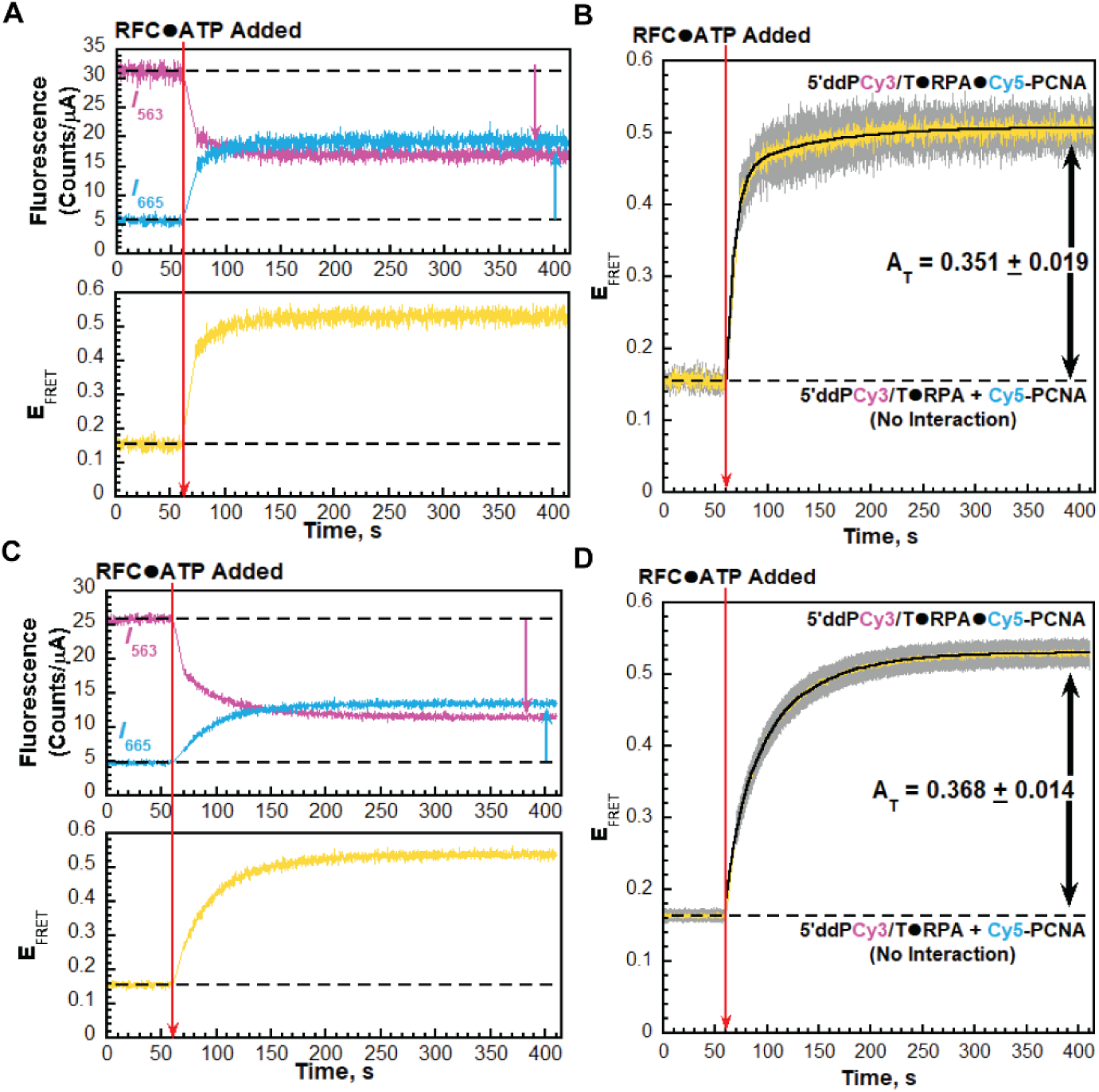
Effect of experimental reaction buffers on RFC-catalyzed loading of PCNA onto P/T junctions. Experiments are carried out on the 5’ddPCy3/T DNA substrate and the results are plotted and analyzed exactly as described in Figure 3A – **C** in the main text. (**A - B**). Data observed with native RPA in Mg^2+^ buffer. (**C – D**) Data observed with native RPA in Ca^2+^/Mg^2+^ buffer. Data is from Figure 3B **– C** in the main text. For E_FRET_ traces in panels **B** and **D**, each is the mean of at least three independent traces with the S.E.M. shown in grey. E_FRET_ traces observed after the addition of RFC are each fit to double exponential rises and the A_T_ are reported in the respective graph as well as in **Table S1** along with other kinetic variables.

**Figure S4.**
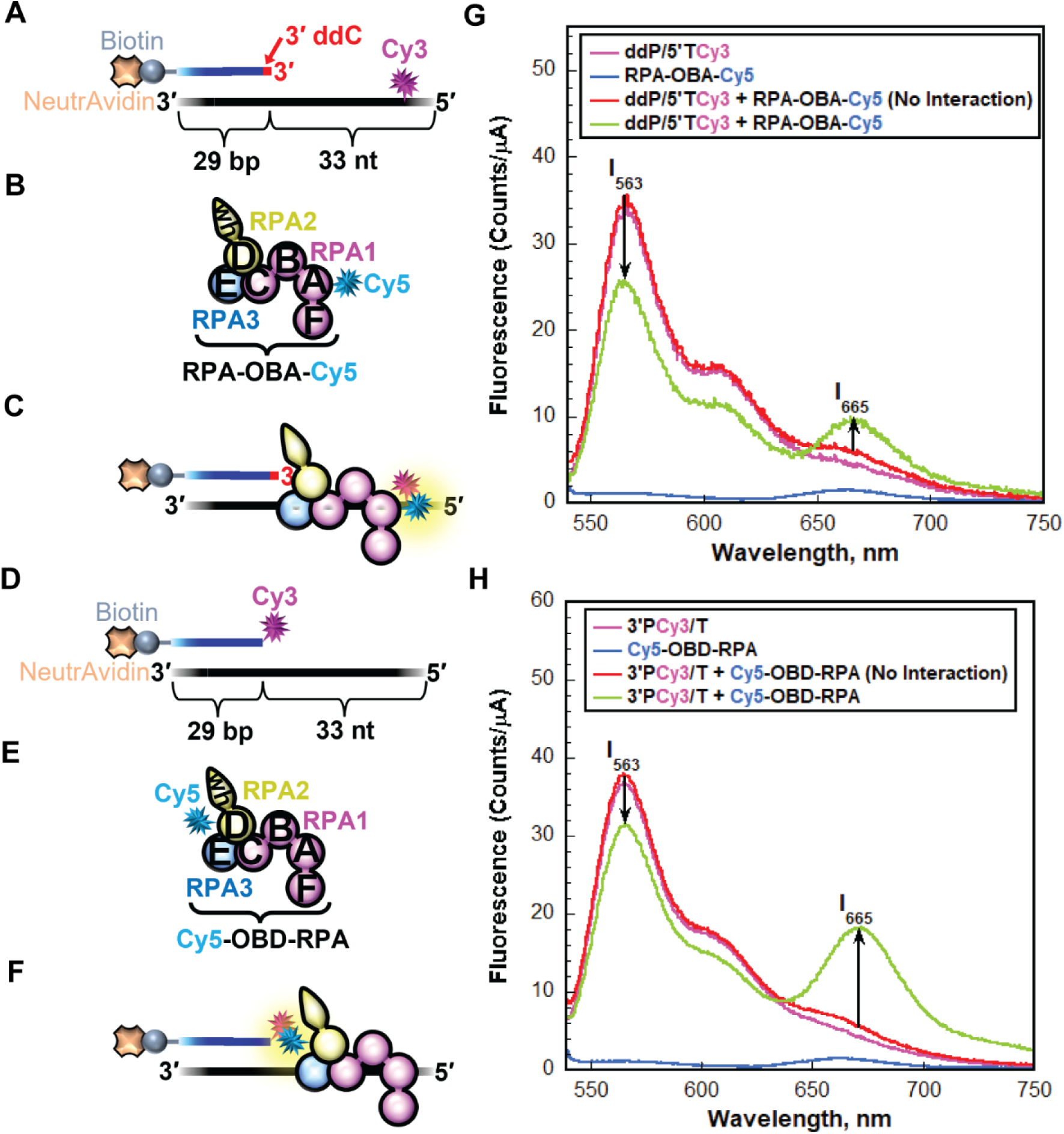
Monitoring interaction between RPA and a P/T junction by FRET. (**A** - **C**) Schematic representations of the Cy3-labeled P/T DNA substrate (panel **A**, ddP/5′TCy3, **Figure S1**) and Cy5-labeled RPA (panel **B**, RPA-OBA-Cy5) utilized to monitor the interaction of RPA OBA with a P/T junction (panel **C**). (**D** – **F**) Schematic representations of the Cy3-labeled P/T DNA substrate (panel **D**, 3′PCy3/T, **Figure S1**) and Cy5-labeled RPA (Cy5-OBD-RPA) utilized to monitor the interaction of OBD with a P/T junction (panel **F**). RPA subunits are color-coded and depicted as in Figure 1. (**G** - **H**) Fluorescence emission spectra of RPA-P/T DNA interactions. *I*_665_ and *I*_563_ are denoted in each spectrum. FRET is indicated by an increase in *I*_665_ and a concomitant decrease in *I*_563_, both of which are denoted in each panel by black arrows. For each panel, the predicted spectrum for no interaction between the respective Cy5-labeled RPA and the corresponding Cy3-labeled P/T DNA is determined by adding the spectrums of the individual components. The spectrums obtained for the interactions between RPA-OBA-Cy5 and the ddP/5′TCy3 DNA substrate and between Cy5-OBD-RPA and the 3′PCy3/T DNA substrate are shown in panels **G** and **H**, respectively.

**Figure S5.**
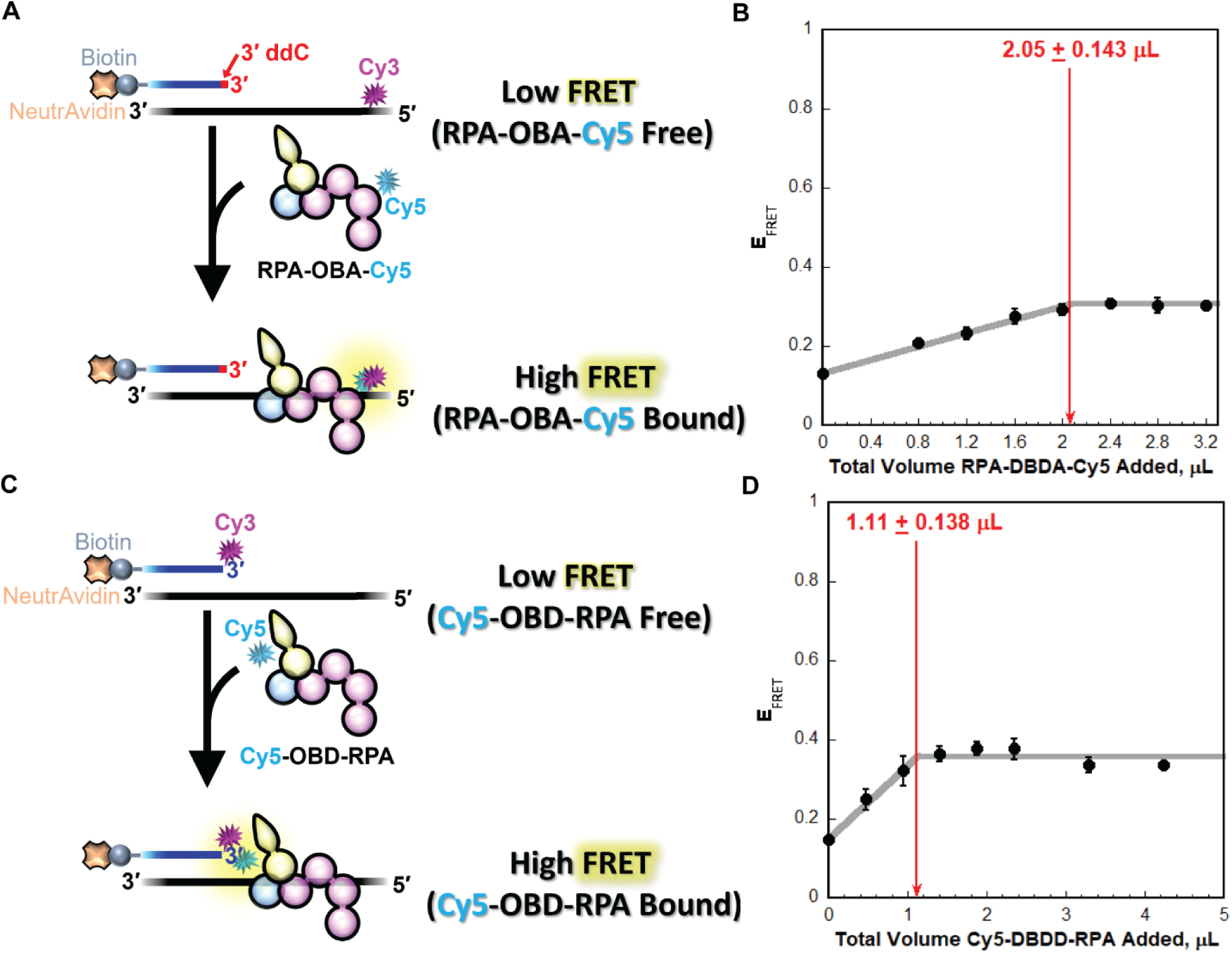
Concentration of active Cy5-labeled RPA. (**A**) Schematic representation of FRET based active site titration of RPA-OBA-Cy5. (**B**) Titration of ddP/5′Cy3T DNA (3.014 pmole, 3.014 pmole RPA binding sites) with RPA-OBA-Cy5. Each data point represents the mean ± S.E.M. of at least three independent measurements. The equivalence point is indicated with standard error of the calculation. Saturation is reached at approximately 2.05 μL total RPA-OBA-Cy5 added, yielding a concentration of 1.47 μM (3.014 pmole RPA binding sites/2.05 μL total RPA-OBA-Cy5 added = 1.47 pmole/μL = 1.47 μM). (**C**) Schematic representation of FRET-based active site titration of Cy5-OBD-RPA. The assay is carried out as described in panel **A** except that the FRET donor is 3′PCy3/T DNA (**Figure S1**) and the FRET acceptor is Cy5-OBD-RPA. (**D**) Titration of 3′PCy3/T DNA (2.43 pmole, 2.43 pmole RPA binding sites) with Cy5-OBD-RPA. Each data point represents the mean ± S.E.M. of at least three independent measurements. The equivalence point is indicated with standard error of the calculation. Saturation is reached at approximately 1.11 μL total Cy5-OBD-RPA, yielding a concentration of 2.19 μM (2.43 pmole RPA binding sites/1.11 μL total Cy5-OBD-RPA added = 2.19 pmole/μL = 2.19 μM).

**Figure S6.**
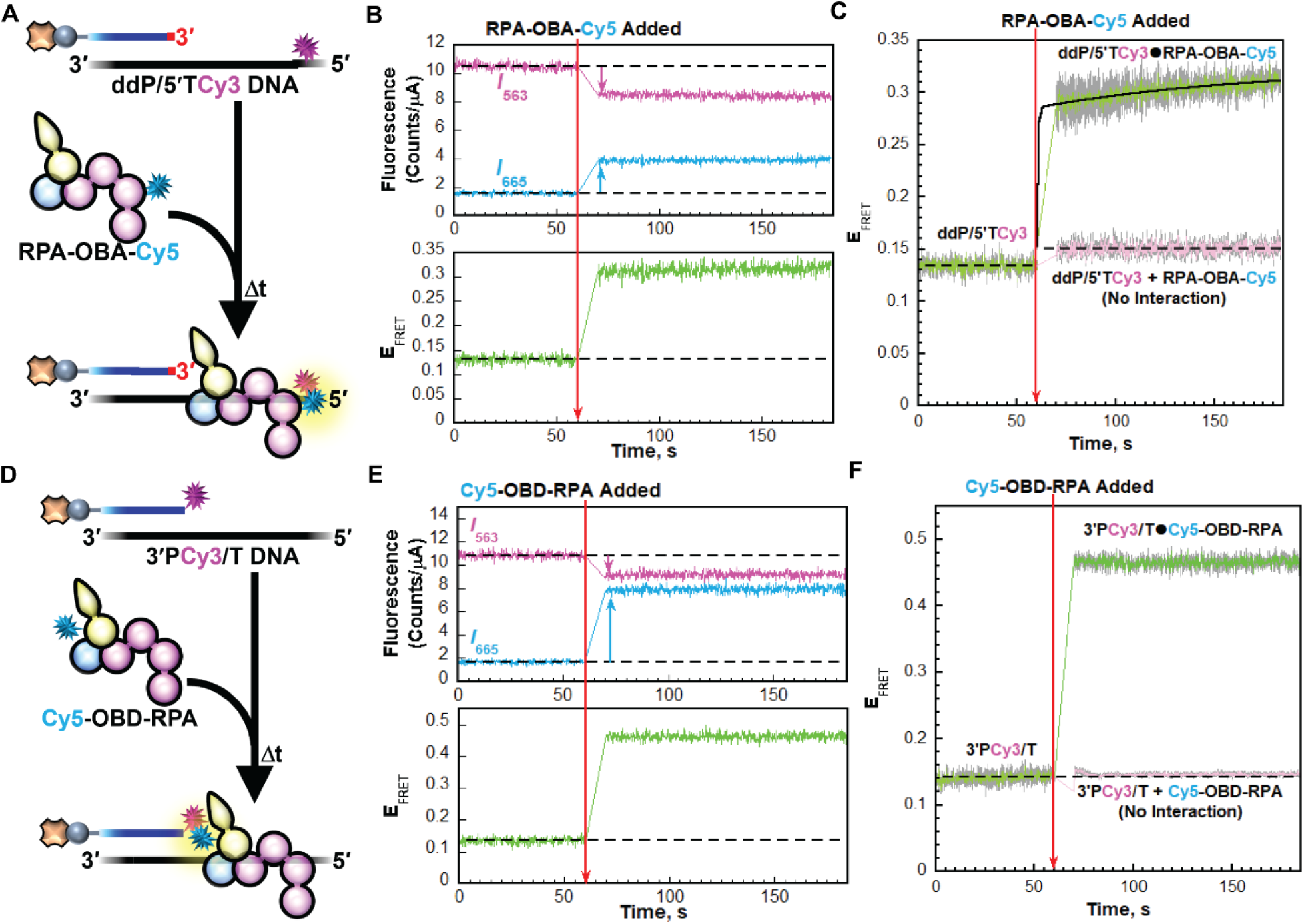
RPA engaging a P/T junction. (**A**) Schematic representation of the FRET experiment performed with ddP/5′TCy3 DNA (+ NeutrAvidin) and RPA-OBA-Cy5 (**B**) Sample time trajectories of *I*_563_ and *I*_665_ (*Top*) and their E_FRET_ (*Bottom*). The time at which RPA-OBA-Cy5 is added is indicated by a red arrow. For observation, the *I*_563_, *I*_665_, and E_FRET_ values observed prior to the addition of RPA-OBA-Cy5 are fit to flat lines that are extrapolated to the axis limits. Changes in *I*_563_ and *I*_665_ are indicated by magenta and cyan arrows, respectively. (**C**) FRET data. Each E_FRET_ trace is the mean of at least three independent traces with the S.E.M. shown in grey. The time at which RPA-OBA-Cy5 is added is indicated by a red arrow. The E_FRET_ trace observed prior to the addition of RPA-OBA-Cy5 is fit to a flat line. The E_FRET_ observed after the addition of RPA-OBA-Cy5 is fit to a double exponential rise and the observed rate constant for the second, slower phase (*k*_obs inc_,_2_) is reported in the graph. The predicted FRET trace (pink) for no interaction between RPA-OBA-Cy5 and the 5′TCy3 DNA is fit to a flat line. (**D** – **F**) Experiments, results, and data analysis carried out for 3′PCy3/T DNA and Cy5-OBD-RPA exactly as described in panels **A** – **C** except FRET traces observed after the addition of Cy5-OBD-RPA were not fit to a kinetic model.

**Figure S7.**
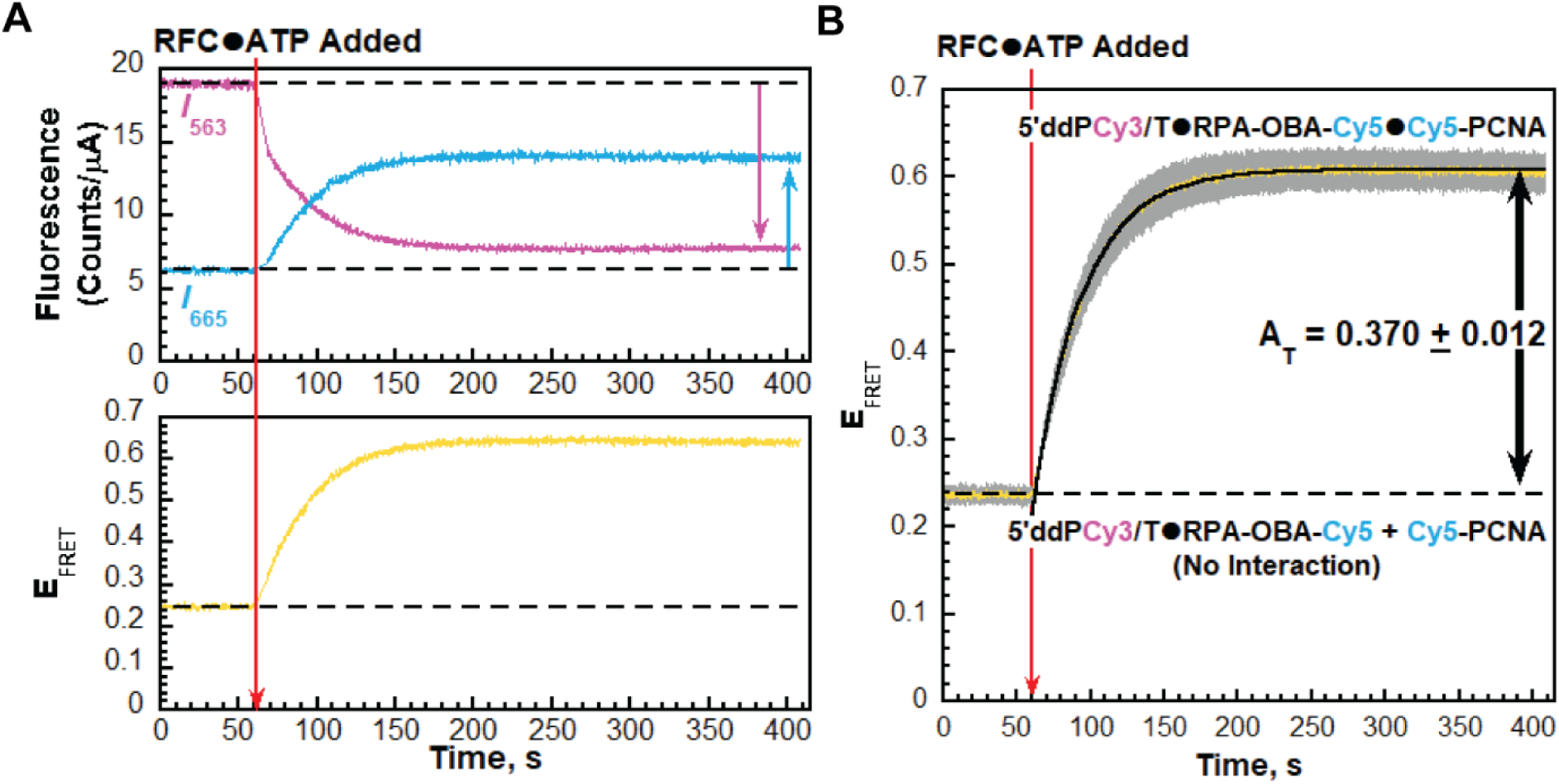
Effect of Cy5 labeling of RPA OBA on RFC-catalyzed loading of PCNA onto P/T junctions. Experiments are carried out with RPA-OBA-Cy5 on the 5′ddPCy3/T DNA substrate and the results are plotted and analyzed exactly as described in Figure 3A – **C** in the main text. E_FRET_ trace in panel **B** is the mean of at least three independent traces with the S.E.M. shown in grey. E_FRET_ trace in panel **B** observed after the addition of RFC is fit to an exponential rise and the A_T_ is reported in the graph as well as in **Table S1** along with other kinetic variables. The total amplitude observed on the 5’ddPCy3/T DNA substrate in Mg^2+^/Ca^2+^ buffer with RPA-OBA-Cy5 (A_T_ = 0.370 ± 0.012) is within experimental error of that observed under the same experimental conditions with native RPA (A_T_ = 0.368 ± 0.014, Figure 3C, **Figure S3D**, **Table S1**). This indicates that RPA-OBA-Cy5 fully supports RFC-catalyzed loading of PCNA onto P/T junctions.

**Figure S8.**
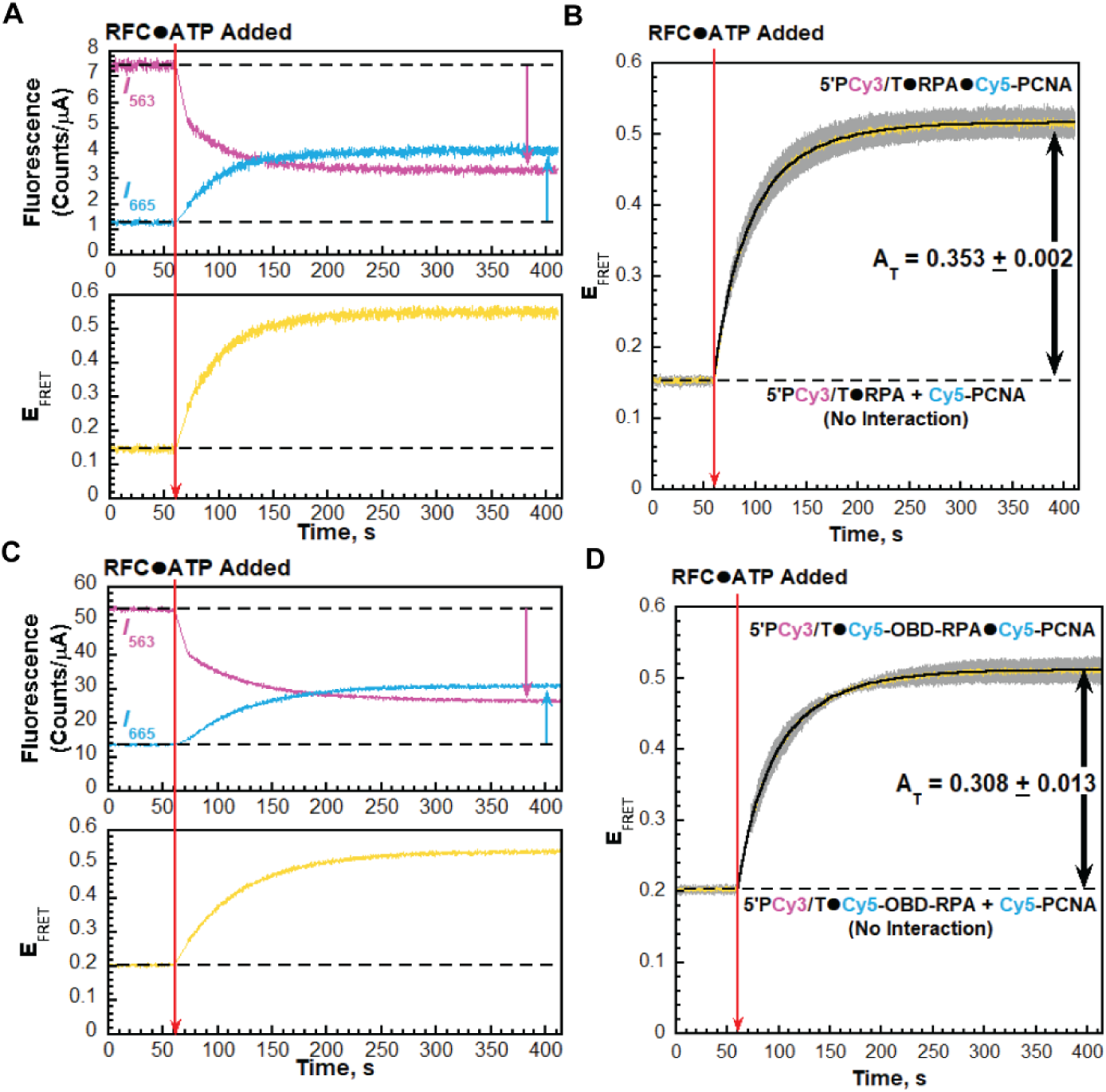
Effect of Cy5 label on OBD of RPA on RFC-catalyzed loading of PCNA onto P/T junctions. Experiments are carried out on the 5′PCy3/T DNA substrate and the results are plotted and analyzed exactly as described in Figure 5A – **C** in the main text. (**A** – **B**) Data observed with native RPA in Ca^2+^/Mg^2+^ buffer (from Figure 5B **– C** in the main text). (**C** – **D**) Data observed with Cy5-ObD-RPA in Ca^2+^/Mg^2+^ buffer. For E_FRET_ traces in panels **B** and **D**, each is the mean of at least three independent traces with the S.E.M. shown in grey. E_FRET_ traces observed after the addition of RFC are each fit to double exponential rises and the total amplitudes (A_T_) are reported in the respective graph as well as in **Table S1** (along with other kinetic variables). The total amplitude observed with Cy5-OBD-RPA (A_T_ = 0.308 ± 0.013) agrees very well with that observed under the same experimental conditions with native RPA (A_T_ = 0.353 ± 0.002, Figure 5C, **Table S1**). Furthermore, the rate constants [*k*_obs inc,1_ = 5.06 ± 0.18 (x 10^-2^) s^-1^, *k*_obs inc,2_ = 1.83 ± 0.02 (x 10^-2^) s^-1^, **Table S1**] observed with Cy5-OBD-RPA are nearly identical to those observed under the same experimental conditions with native RPA [*k*_obs inc,1_ = 4.34 ± 0.18 (x 10^-2^) s^-1^, *k*_obs inc,2_ = 1.79 ± 0.04 (x 10^-2^) s^-1^, **Table S1**]. Together, this indicates that RPA-OBA-Cy5 fully supports RFC-catalyzed loading of PCNA onto P/T junctions.

**Figure S9.**
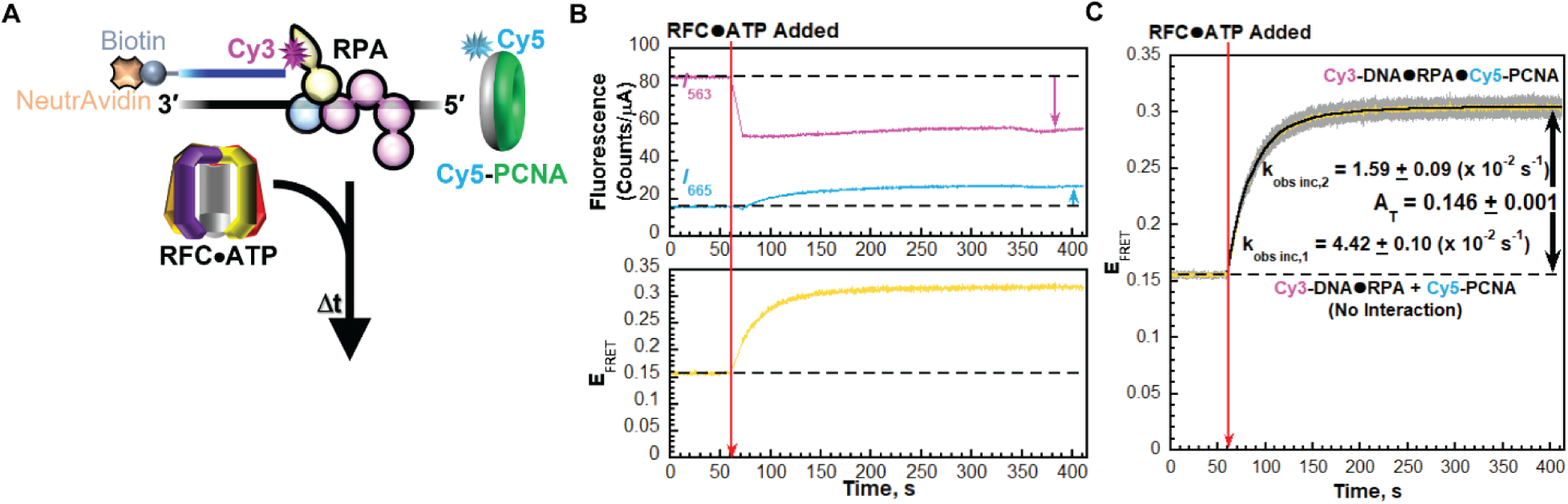
Effects of primer Cy3 donor location on the kinetics RFC-catalyzed loading of PCNA onto P/T junctions. (**A**) Schematic representation of the FRET experiment. Reactions were carried out exactly as described in **5A** in the main text except with 3′Cy3P/T DNA. Note the Cy5 label on PCNA will be oriented away from the Cy3 donor on the 3′ terminus of the primer strand when Cy5-PCNA is loaded onto the 3′Cy3P/T DNA substrate by RFC (**B**) Sample time trajectories of *I*_563_ and *I*_665_ (*Top*) and their E_FRET_ (*Bottom*). The time at which the RFC•ATP complex is added is indicated by a red arrow. Changes in *I*_563_ and *I*_665_ are indicated by magenta and cyan arrows, respectively. For observation, the *I*_563_, *I*_665_, and E_FRET_ values observed prior to the addition of the RFC•ATP complex are fit to flat lines that are extrapolated to the axis limits. (**C**) FRET data. Each E_FRET_ trace is the mean of at least three independent traces with the S.E.M. shown in grey. The time at which the RFC•ATP is added is indicated by a red arrow. The E_FRET_ trace observed prior to the addition of the RFC•ATP complex represents the complete absence of interactions between the 3′PCy3/T•RPA complex and Cy5-PCNA and is fit to a flat line that is extrapolated to the axis limits. The E_FRET_ trace observed after the addition of the RFC•ATP complex is fit to a double exponential rise and the observed rate constants (*k*_obs inc,1_ and *k*_obs inc,2_) and A_T_ are reported in the graph as well as in **Table S1**. The predicted E_FRET_ trace (pink) for no interaction between 5P′Cy3/T•RPA complex and the loading complex is fit to a flat line.

**Figure S10.**
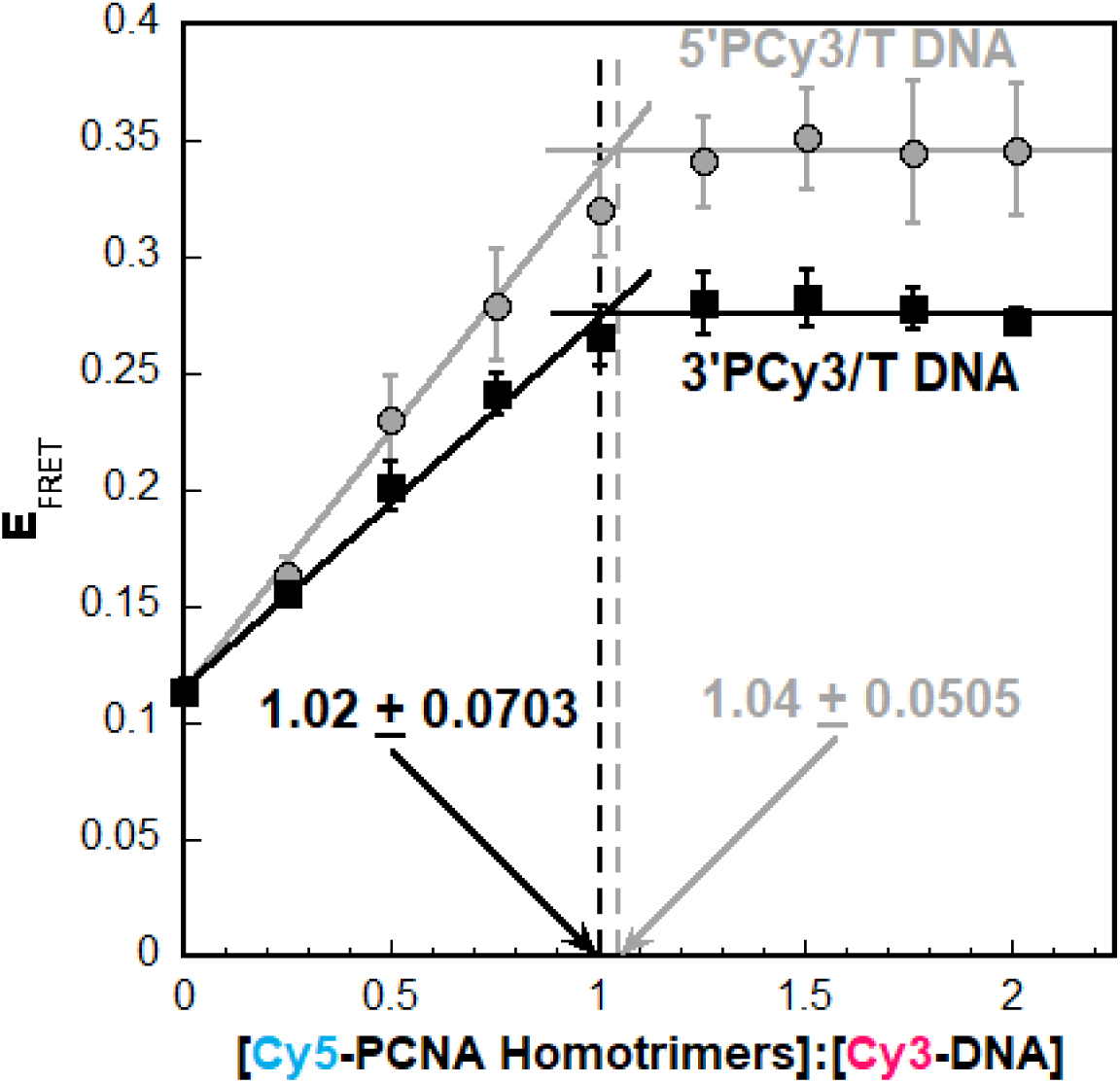
Titrations of the steady state FRET signal. Results are plotted as a function of the [Cy5-PCNA homotrimer]:[Cy3-DNA] ratio and each data point represents the mean ± S.E.M. of at least three independent measurements. Data observed for the 5′PCy3/T and 3′PCy3/T DNA substrates is shown in grey and black, respectively. Under the experimental conditions, RFC-catalyzed loading of PCNA onto the 5′PCy3/T is stoichiometric (**Figure S9**) and, hence, FRET increases linearly until the DNA is saturated with PCNA (i.e., the equivalence point)^2,3^ at a ratio of 1 PCNA homotrimer:1 P/T DNA. Data is fit to two segment lines (a linear regression with a positive slope and a flat line) and the equivalence points (indicated with standard errors of the calculations) are calculated from the intersection of the two segment lines. Equivalence points for each P/T DNA substrate are indicated on the graph.

**Figure S11.**
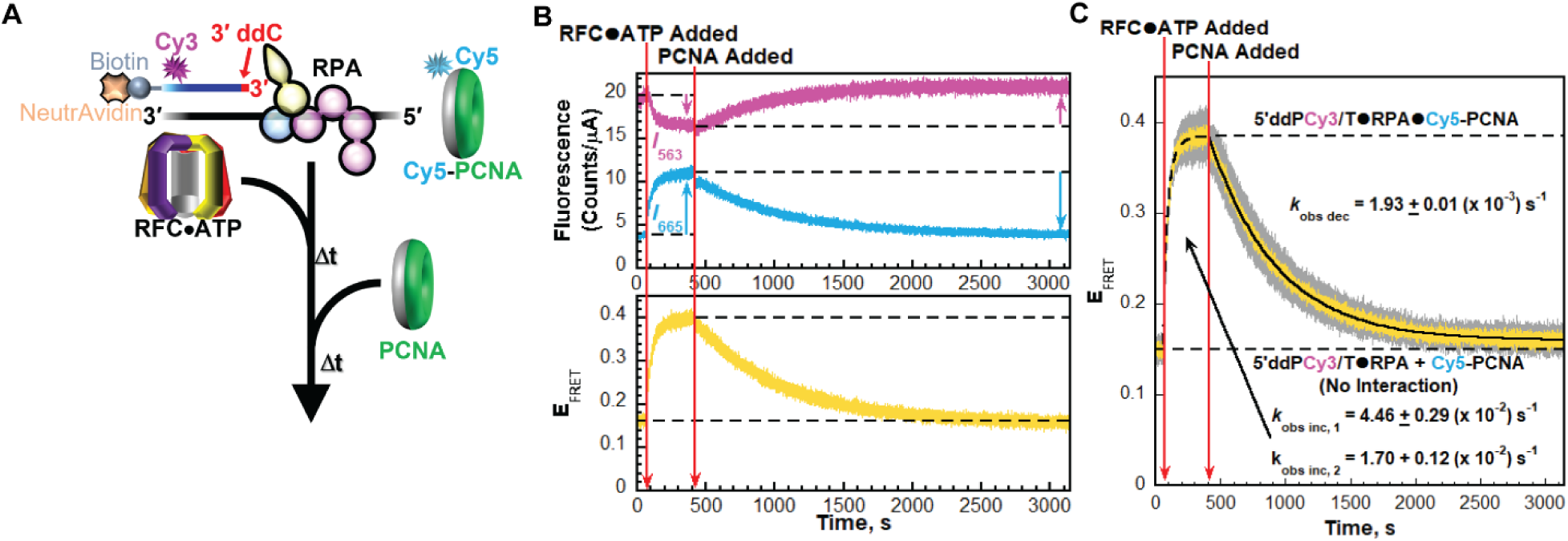
Dynamics of PCNA encircling a P/T junction. (**A**) Schematic representation of the FRET experiment performed with 5′ddPCy3/T DNA, RPA, Cy5-PCNA, RFC, ATP, and PCNA. (**B**) Sample time trajectories of *I*_563_ and *I*_665_ (*Top*) and their E_FRET_ (*Bottom*). The times at which the RFC•ATP complex and unlabeled PCNA are added are indicated by red arrows. For observation, the emission intensity traces and E_FRET_ values observed in the absence of RFC are each fit to dashed flat lines that are extrapolated. Also, dashed flat lines are drawn to highlight the *I* values and E_FRET_ values observed at equilibrium after the addition of unlabeled PCNA. Changes in *I*_563_ and *I*_665_ observed after each addition are indicated by magenta and cyan arrows, respectively. (**C**) FRET data. Each E_FRET_ trace is the mean of at least three independent traces with the S.E.M. shown in grey. The times at which RFC•ATP and unlabeled PCNA are added are indicated by red arrows. The E_FRET_ trace observed prior to the addition of the RFC•ATP complex is fit to a dashed flat line that is extrapolated to the axis limits to depict the average E_FRET_ value for no interaction between Cy5-PCNA and the 5′ddPCy3/T•RPA complex. The E_FRET_ trace observed after the addition of the RFC•ATP complex is fit to a dashed double exponential rise that is extrapolated to the axis limits to depict the average E_FRET_ value for complete loading of Cy5-PCNA onto the Cy3-labeled P/T DNA substrate (i.e., the 5′ddPCy3/T•RPA•Cy5-PCNA complex). The rate constants observed for the E_FRET_ increase (*k*_obs inc, 1_ and *k*_obs inc, 2_) are reported in the figure. The E_FRET_ trace observed after the addition of unlabeled PCNA is fit to a single exponential decay and the observed rate constant (*k*_obs dec_) is reported in the graph.

**Figure S12.**
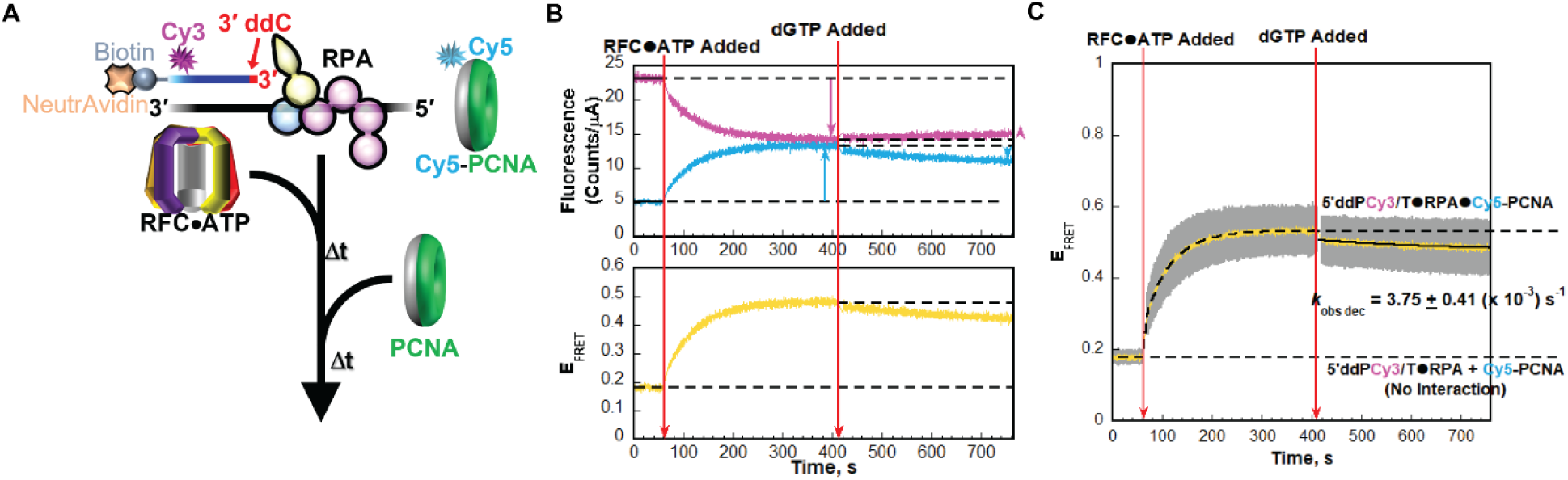
Effects of added dGTP on the dynamics of PCNA encircling a P/T junction. (**A**) Schematic representation of the FRET experiment performed with 5′ddPCy3/T DNA, RPA, Cy5-PCNA, RFC, ATP, and dGTP. (**B**) Sample time trajectories of *I*_563_ and *I*_665_ (*Top*) and their E_FRET_ (*Bottom*). The times at which the RFC•ATP complex and dGTP are added are indicated by red arrows. For observation, the emission intensity traces and E_FRET_ values observed in the absence of RFC are each fit to dashed flat lines that are extrapolated. Also, dashed flat lines are drawn to highlight the *I* values and E_FRET_ values observed at equilibrium after the addition of RFC•ATP. Changes in *I*_563_ and *I*_665_ observed after each addition are indicated by magenta and cyan arrows, respectively. (**C**) FRET data. Each E_FRET_ trace is the mean of at least three independent traces with the S.E.M. shown in grey. The times at which RFC•ATP and dGTP are added are indicated by red arrows. The E_FRET_ trace observed prior to the addition of the RFC•ATP complex is fit to a dashed flat line that is extrapolated to the axis limits to depict the average E_FRET_ value for no interaction between Cy5-PCNA and the 5′ddPCy3/T•RPA complex. The E_FRET_ trace observed after the addition of the RFC•ATP complex is fit to a dashed double exponential rise that is extrapolated to the axis limits to depict the average E_FRET_ value for complete loading of Cy5-PCNA onto the Cy3-labeled P/T DNA substrate (i.e., the 5′ddPCy3/T•RPA•Cy5-PCNA complex). The E_FRET_ trace observed after the addition of dGTP is fit to a single exponential decay and the observed rate constant (*k*_obs dec_) is reported in the graph.

**Figure S13.**
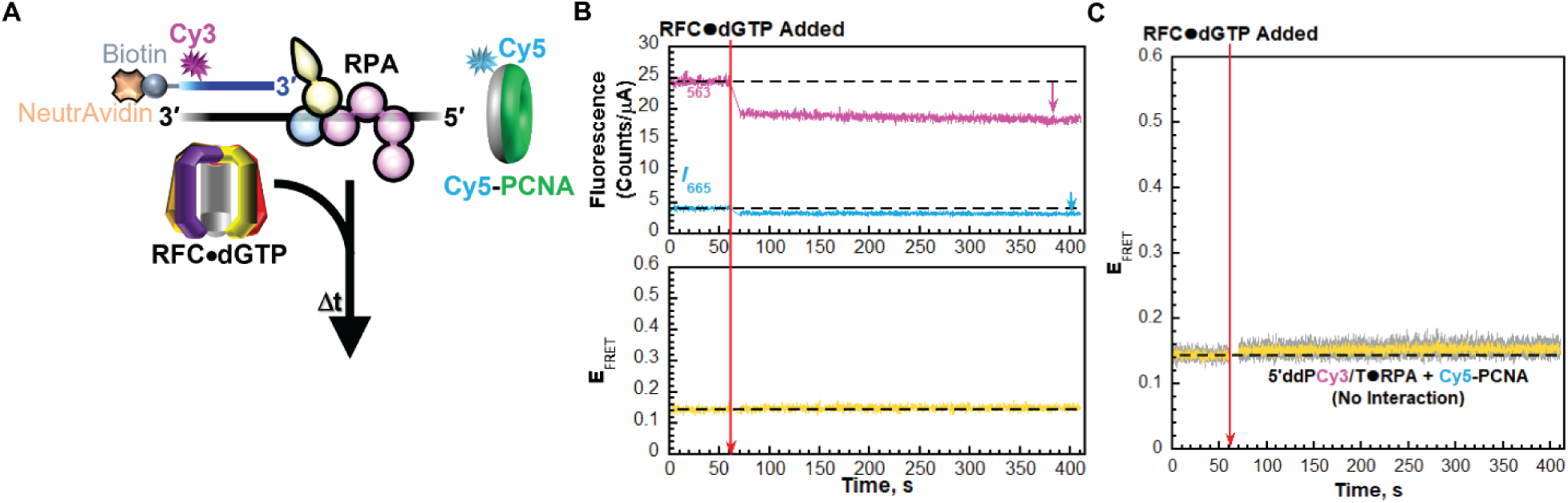
Effects of dGTP on RFC-catalyzed loading of PCNA onto a P/T junction. (**A**) Schematic representation of the FRET experiment performed with 5′ddPCy3/T DNA, RPA, Cy5-PCNA, RFC, and dGTP. (**B**) Sample time trajectories of *I*_563_ and *I*_665_ (*Top*) and their E_FRET_ (*Bottom*). The time at which the RFC•dGTP complex is added is indicated by a red arrow. For observation, the emission intensity traces and E_FRET_ values observed in the absence of RFC are each fit to dashed flat lines that are extrapolated. Changes in *I*_563_ and *I*_665_ observed after each addition are indicated by magenta and cyan arrows, respectively. (**C**) FRET data. Each E_FRET_ trace is the mean of at least three independent traces with the S.E.M. shown in grey. The times at which RFC•dGTP is added is indicated by red arrows. The E_FRET_ trace observed prior to the addition of the RFC•dGTP complex is fit to a dashed flat line that is extrapolated to the axis limits to depict the average E_FRET_ value for no interaction between Cy5-PCNA and the 5′ddPCy3/T•RPA complex.

**Figure S14.**
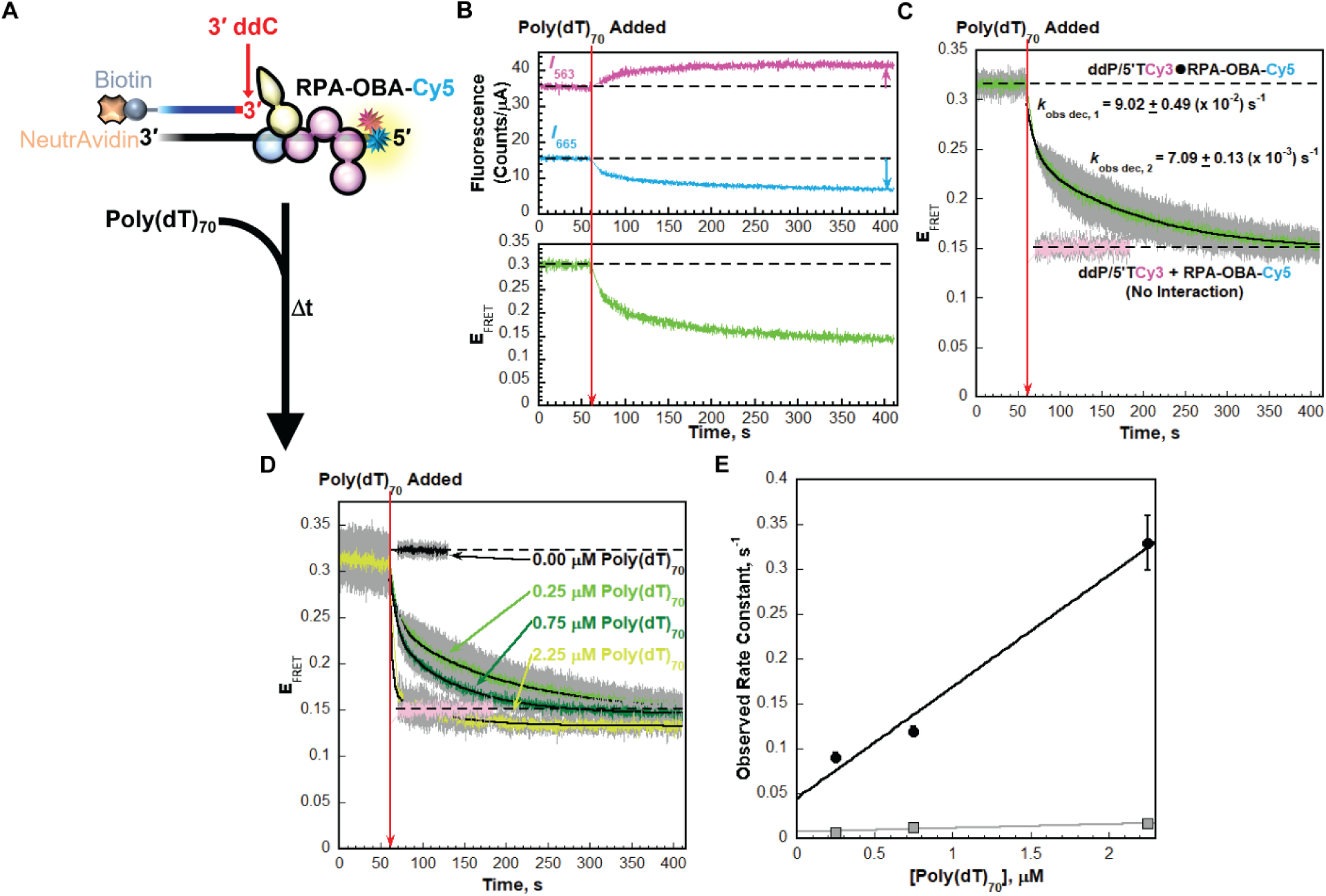
Facilitated exchange of RPA between ssDNA sequences. (**A**) Schematic representation of the FRET experiment performed with ddP/5′TCy3 DNA, RPA-OBA-Cy5, ATP, and poly(dT)_70_. (**B**) Sample time trajectories of *I*_563_ and *I*_665_ (*Top*) and their E_FRET_ (*Bottom*) observed with 0.235 μM poly(dT)_70_. The time at which poly(dT)_70_ is added is indicated by a red arrow. For observation, the emission intensity traces and E_FRET_ values observed in the absence poly(dT)_70_ are each fit to dashed flat lines that are extrapolated. Changes in *I*_563_ and *I*_665_ observed after each addition are indicated by magenta and cyan arrows, respectively. (**C**) FRET data observed in the presence of 0.250 μM poly(dT)_70_. Each E_FRET_ trace is the mean of three independent traces with the S.E.M. shown in grey. The times at which poly(dT)_70_ is added is indicated by a red arrow. The E_FRET_ trace observed prior to the addition of poly(dT)_70_ is fit to a dashed flat line that is extrapolated to the axis limits. The E_FRET_ trace observed after the addition of poly(dT)_70_ is fit to a double exponential decline and the observed rate constants (*k*_obs dec,1_ and *k*_obs dec,2_) are reported in the graph. The predicted E_FRET_ trace (pink) for no interaction between RPA-OBA-Cy5 and the ddP/5′TCy3 DNA is fit to a flat line. (**D**) FRET data observed in the presence of increasing concentrations of poly(dT)_70_. Each E_FRET_ trace is the mean of three independent traces with the S.E.M. shown in grey. The times at which poly(dT)_70_ is added is indicated by a red arrow. The E_FRET_ trace observed after the addition of buffer is fit to a flat line. The E_FRET_ traces observed after the addition of a non-zero concentration of poly(dT)_70_ are fit to double exponential declines and the observed rate constants (*k*_obs dec,1_ and *k*_obs dec,2_) are plotted in panel **E** as a function of poly(dT)_70_ concentration.

**Table S1.**
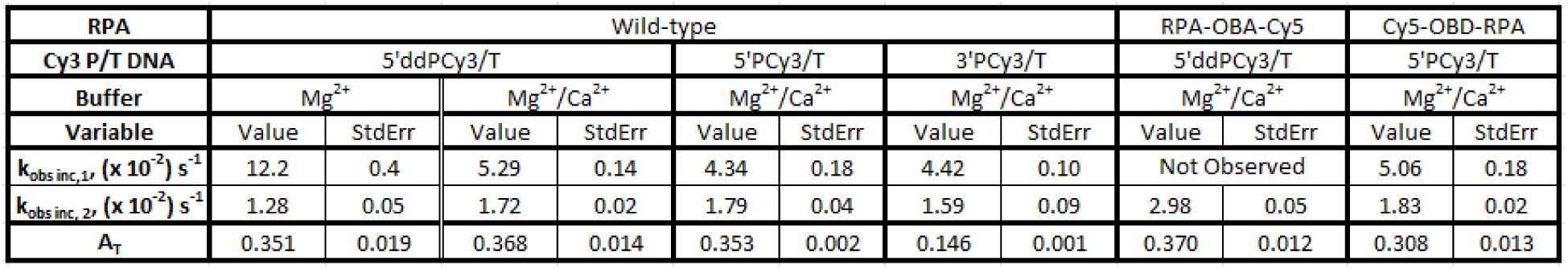
Kinetic variables of RFC-catalyzed loading of PCNA onto a P/T junction.

## Footnotes

1 The content is solely the responsibility of the authors and does not necessarily represent the official views of the National Institutes of Health

## Notes

### Competing Interest Statement

The authors have declared no competing interest.

